# Silent hippocampal engrams shape future memory processing

**DOI:** 10.64898/2026.04.12.717978

**Authors:** Ali Choucry, Kareem Abdou, Kaoru Inokuchi

**Affiliations:** Research Centre for Idling Brain Science, University of Toyama, Toyama 930-0194, Japan; Department of Biochemistry, Graduate School of Medicine and Pharmaceutical Sciences. University of Toyama, Toyama, Japan; CREST, Japan Science and Technology Agency (JST), University of Toyama, Toyama, Japan; Department of Pharmacology and Toxicology, Faculty of Pharmacy, Cairo University, Cairo 11562, Egypt; Department of Biochemistry, Faculty of Pharmacy, Cairo University, Cairo, Egypt; College of Pharmacy, Al-Ain University, Abu Dhabi, UAE

**Author notes:** To whom correspondence should be addressed: Kaoru Inokuchi.

## Abstract

The hippocampus harbors silent memory traces even when memory recall is absent. It is unknown why these traces persist and whether they remain sensitive to future episodes. We trained mice on the hippocampal-dependent novel object place recognition task and tracked the transition of the engram from a latent to a silent phase, which still held a functional memory trace. When mice were exposed to a weaker, modified version of their original memories, silent engrams led to better consolidation of these new events than their latent predecessors. Bidirectional manipulation of the engram’s state, as well as artificially prolonging its latent state, successfully reversed its effect on the second memory. The silent engram state consistently displayed higher neuronal reactivation during the second event compared to the latent one, which may explain its preferential effect on its consolidation. Our results reveal previously unknown cognitive and activation dynamics for this behaviorally silent state.

## Introduction

Memories are allocated to a group of neurons termed an engram, which is both necessary^1^ and sufficient^2^ for memory recall. The hippocampus encodes high-level representations of episodic memories, weaving together specifics into a detailed, yet labile trace^3,4^. This short-term memory may next be stabilized through a consolidation process, forming a stable, long-term memory^5,6^. Finally, systems consolidation transfers this detailed hippocampal event memory into a cortical abstract representation, which is maintained and used when recall is invoked at remote time points^7,8^, a process where engram cells are dynamically involved^9–12^. This linear, overly simplified narrative has recently been challenged with multiple instances of a more nuanced memory storage and recall process. For example, short-term memories are maintained in the hippocampus for long-term intervals^13^. Furthermore, long-term memories whose consolidation was disrupted can still be expressed by artificially stimulating their engram hosts^14^, even days later^15^. Different engram states thus exist and dictate the means for successful memory recall. An “active” state denotes an engram that is currently retrieved, a “latent” engram activates spontaneously without any artificial manipulation, and a “silent” engram describes a state that responds to artificial stimulation but not to natural cues^16^. Interestingly, silent hippocampal engrams exist both with^12^ and without^13–15,17^ spontaneous memory recall, and express weaker reactivation levels compared to their latent predecessors. In the former case, hippocampal-cortical dialogue necessary for systems consolidation is thought to silence the hippocampal engram once a cortical trace has matured enough to support recall. In the absence of such cortical recall, silent hippocampal traces show weakened spines but maintain their downstream engram connectivity. Interestingly, they also display a graded behavioral response that scales with the strength of their activation^15^, hinting at a thresholded, rather than binarized, readout of hippocampal output. Moreover, silent hippocampal engrams can spontaneously revert to a latent-like state following exposure to a reminder cue^17,18^ or a social experience^19^. Similarly, maturation of hippocampal engrams from latent to silent could be halted^17,20^ or accelerated^17^ by various manipulations.

Among the chief modulators of neuronal plasticity^21^ that prolong the latent state of hippocampal engrams^20^ are the calcium-impermeable (GluA_2_-containing) α-amino-3-hydroxy-5-methyl-4-isoxazolepropionic acid receptors (AMPARs). Synaptic GluA_2_-containing AMPARs correlate with the strength and maintenance of long-term memories^22,23^, and enhance the retrievability of silent hippocampal engrams^20,24^. Synthetic decoy peptides such as GluA23Y mimic key regions required for the internalization of these receptors, keeping them at the synaptic surface ^22,24–28^.

Another departure from the linear engram allocation/reactivation/memory recall pathway is the mounting experimental and clinical evidence for the lasting effects of forgotten memories. From altering gene expression in *C. elegans*^29^ to reinstating encoding activity^30^ and guiding human behavior in various associative memory tasks^31–35^, silent memory traces can evidently yet covertly still be engaged, sometimes even a decade after their inception^33^.

Taking what we know about silent engrams, it may seem counterintuitive to maintain memory traces without a reliable spontaneous recall until we consider the possibility that silent engrams may still be providing more covert function(s). Indeed, forgetting is no longer considered a failure of a cognitive system, but rather a necessary sacrifice of specifics, away from over-fitting, a dynamic process of adaptive learning^36^. The hippocampus actively participates in rapid, higher-order episodic management, extracting regularities^37–42^, building mnemonic structures^43,44^, and both integrating and separating learned information within the same behavioral epoch^45^. Indeed, an elegant neural network model demonstrated that the hippocampus can process both regularities and specifics across episodes^46^. Experimentally, significant hippocampal engram overlap occurs with related events^37,47^, mirroring overlaps observed in extra-hippocampal^48^ and cortical^49^ regions, highlighting its ability to compute higher-order relationships among episodes^50^. Collectively, compelling computational, experimental and clinical evidence confirms an active hippocampal role beyond simple pattern separation, including a more expansive evaluation of different events with common and unique features. The aforementioned literature expanded our understanding of hippocampal silent engrams, but is limited by several key design choices. Some studies, for example, artificially enforced the silent engram state through interfering with memory consolidation^14,15^. In studies where the engram was allowed to consolidate normally, systems consolidation still supported natural memory recall^10,12^. When natural forgetting was combined with the lack of spontaneous recall, silent hippocampal engrams were primarily studied within the scope of their original memories^17^. These studies, therefore, do not address why the hippocampus maintains silent engrams of naturally consolidated memories instead of making them immediately unavailable when forgetting occurs.

In this study, we investigated how these silent engrams behave when they encounter a weak, modified episode that was contextually linked to their original event. We found that, while both latent and silent engrams allowed the formation of short-term memories of the new event, only the silent state processed the weak event into a stronger long-term memory. Artificially inhibiting or activating the latent and silent states reversed their outcomes on the new event. Both the engram state transition and the subsequent cognitive outcome were also sensitive to the synaptic localization of GluA_2_-containing AMPARs. Immunohistochemical assessment revealed that a higher proportion of the silent engram was reactivated when encountering the modified event compared to its latent counterpart. Our data thus support a model where the hippocampus maintains silent engram traces that can still be significantly reactivated by future events for memory generalization.

## Results

### 1. Natural timeline for state transition of the hippocampal object-place memory engram

We chose the novel object place recognition (NOPR) task for its hippocampus-dependent memory^51^, which is naturally forgotten over time^20^. Mice were placed in a square context with two identical copies of an object in two opposite corners. After a single 15-minute encoding session, mice were tested one to five days later, with one object moved to a novel location (Fig.1a and Extended Data Fig.1a, b). Object location memory was well preserved one and two days after encoding, as mice spent more time exploring the novel location object than the stationary one. At three and five days, however, the overall performance dropped to non-significant levels, insinuating either weakening or a complete deletion of the memory. To confirm the presence of a silent memory trace, we labelled the NOPR training engram with a stimulatory chemo-genetic actuator, and five days later, we activated it with clozapine-N-oxide (CNO) (or its vehicle (VEH) as a control) before the test (Fig.1b, c and Extended Data Fig.1c, d). This stimulation successfully rescued the NOPR memory, indicating that two days after encoding, the engram naturally shifts from a latent to a silent state, which is maintained for at least three more days, in agreement with previous reports^15^. The same approach could not induce significant discrimination at a more remote time point (Fig.1d and Extended Data Fig.1e, f). These results outline the time course by which the hippocampal NOPR engram state naturally shifts from latent to silent and then to unavailable.

**Figure 1:**
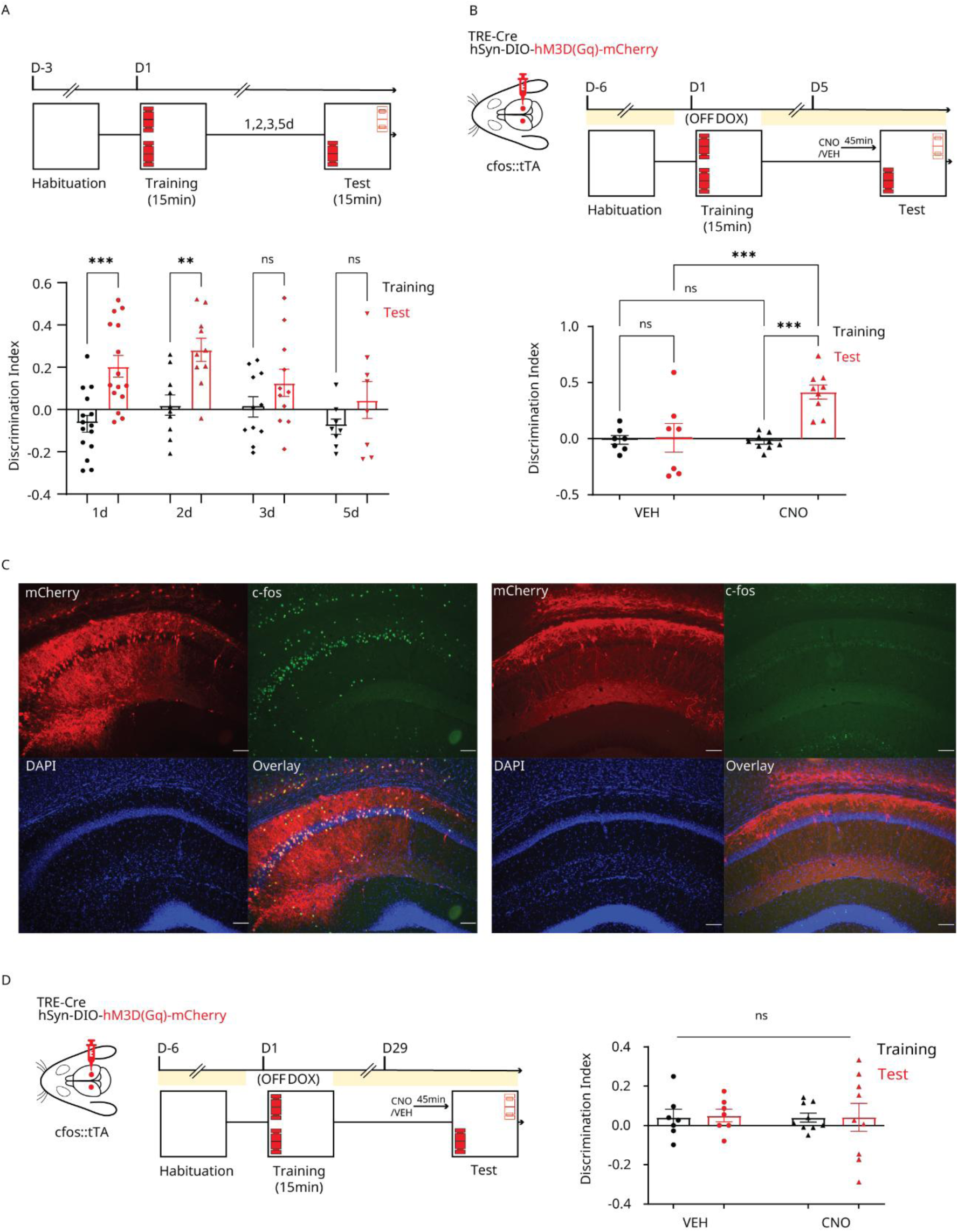
Silent NOPR engram. **(A)** Top, behavioral timeline to check for object-place memory persistence. Bottom: discrimination of the novel object’s location during training and testing at different intervals. *n* = 15, 10, 11, and 8 mice for 1, 2, 3, and 5 days, respectively. **(B)** Top, design and behavioral timeline of the chemo-genetic memory rescue experiment. Bottom, discrimination of the novel object’s location during training and testing at 5 days. *n* = 7 and 9 mice for VEH and CNO, respectively. **(C)** Labelling and CNO (left) or VEH (right) activation of engram cells in CA1. Brain sections were immunostained for mCherry (red), c-Fos (green) and DAPI (blue). Scale bars, 100 μm. **(D)** Left, design and behavioral timeline of the remote chemo-genetic memory rescue experiment, similar to **(B)** except that testing was done after 28 days. Right, discrimination of the novel object’s location during training and testing at 28 days. *n* = 7 and 9 mice for VEH and CNO, respectively. All statistical comparisons were performed using repeated measures two-way ANOVA. CNO, clozapine-N-oxide; VEH, vehicle. ns, non-significant; **, *P* < 0.01; ***, *P* < 0.001. Data are represented as mean ± standard error of the mean (SEM).

### 2. Different outcomes of latent and silent engrams in a dual NOPR task

We next tested whether latent and silent NOPR engrams react differently to a modified NOPR event of high conceptual similarity. First, we confirmed that a single 4-minute NOPR session failed to elicit significant long-term object location memory (Extended Data Fig.2), in agreement with previous reports^52^. To establish a dual NOPR paradigm, we changed the flooring and wall texture of the original context and attached unique patterns to create a modified iteration that shares the same geometric properties but provides distinct tactile and visual features. For four consecutive days, we habituated mice to both iterations, with a 10-minute session in each context separated by 30 minutes. This habituation schedule aimed to enforce a link between contexts through the established temporal window for prospective linking^37,47,50^ as well as through repeated co-exposure^48^. Mice were then trained in the original context with a set of objects for 15 minutes (the original event). A modified event consisted of a 4-minute session in the modified context with a novel set of objects placed in the same corners as the original event, regardless of the locations of contextual cues (Extended Data Fig.3a-c). Mice were exposed to this modified event either one day (the latent group) or four days (the silent group) after the original event, and were tested on the following day (Fig.2a and Extended Data Fig.3). We also checked the original event’s memory for both groups 5 days after its encoding, when the NOPR recall should be absent (Fig.1).

**Figure 2:**
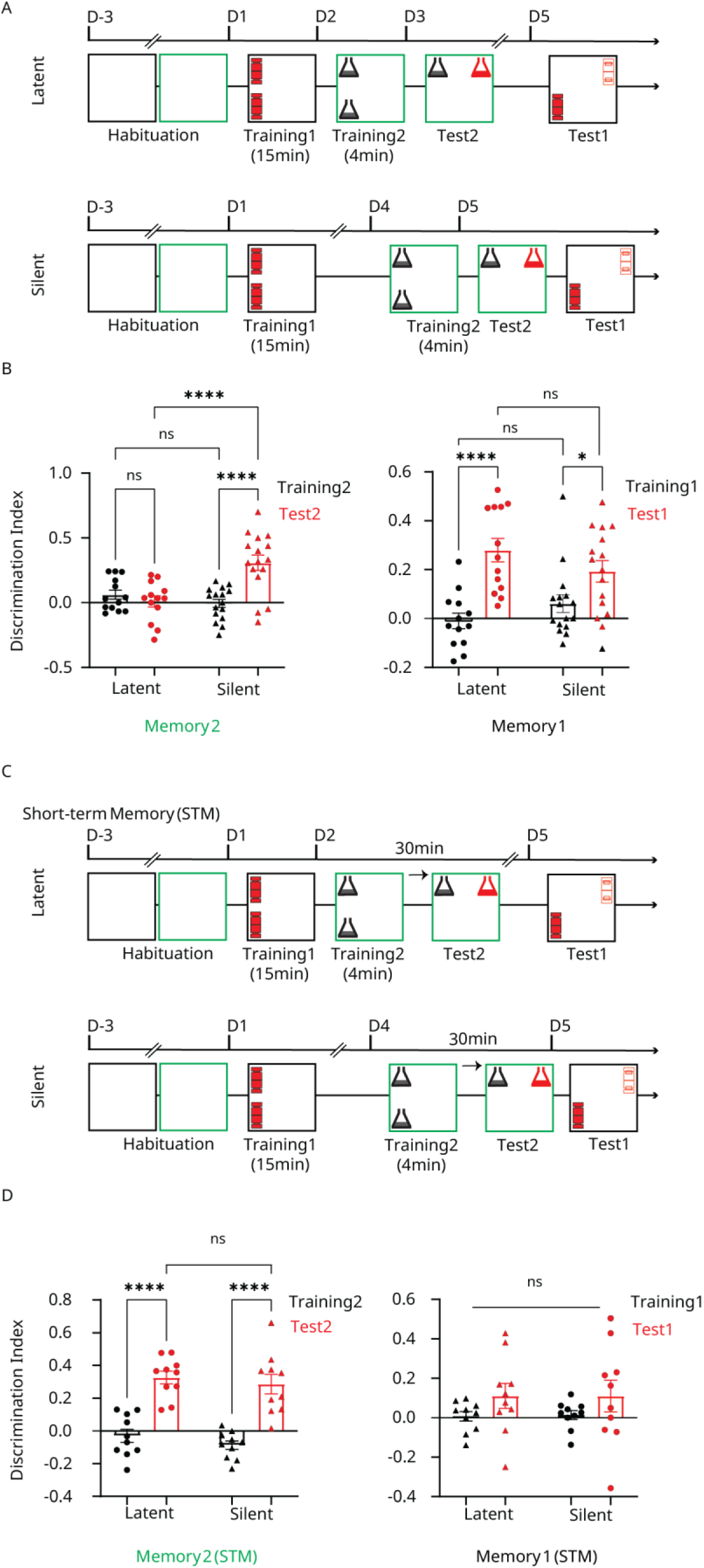
Dual NOPR paradigm with latent and silent engrams. **(A)** Behavioral timeline for the latent (top) and silent (bottom) NOPR engrams. **(B)** Discrimination of the novel object’s location at the modified (left) and original (right) memory test for the latent and silent groups. *n* = 13 and 16 mice for the latent and silent groups, respectively. **(C) and (D)** Same as **(A) and (B),** but for the short-term memory test. *n* = 10 mice/group. All statistical comparisons were performed using repeated measures two-way ANOVA. ns, non-significant; *, *P* < 0.05; ****, *P* < 0.0001. Data are represented as mean ± standard error of the mean (SEM).

In the modified memory test, the latent group showed no discrimination of the novel object location, as opposed to when the modified event was encoded during the silent state, where mice strongly preferred the moved object (Fig.2b). These differences were not due to discrepancies in the modified training (Extended Data Fig.3d, e). The modified NOPR event also induced a spontaneous reversal of the original engram into a latent state, as both groups recovered their original memories (Fig.2b and Extended Data Fig.3f, g). We repeated this experiment using a modified schedule with a longer temporal gap between latent and silent encoding. Instead of the two-day difference in the previous experiment, mice were exposed to the modified event three hours or five days after encoding the original NOPR memory for the latent and silent groups, respectively (Extended Data Fig.4a). Again, both groups recovered their original memories, while only the latent group showed significant discrimination at the modified memory test (Extended Data Fig.4b-f). This experiment further supports our claim that the behavioral discrepancies between our groups are induced by discrete engram states as opposed to other, non-specific temporal artifacts, as we utilized the earliest time point post consolidation sleep^53^ (where the latent engram should be at its strongest), and a more delayed time point for the silent engram where we repeatedly confirmed its lack of spontaneous recall (Fig.1a, b). Our results so far, however, do not explain whether the state of the original NOPR engram affects the encoding or the consolidation of the modified iteration. To disentangle these possibilities, we evaluated the short-term memory of the modified event in a new cohort of mice that underwent the same dual paradigm (Fig.2c, d and Extended Data Fig.3h-k). Both groups successfully discriminated the novel location-object in the modified memory test, showing that learning was not affected by the current engram state. The original memory, however, failed to recover significantly in both groups. Collectively, our dual NOPR paradigm employed two distinct yet spatially similar events in repeatedly co-exposed contexts. Both the latent and silent engrams learned the new event equally well, but only the silent state improved its consolidation.

### 3. Bidirectional manipulation of engram state before the modified event

Many studies investigated the dynamics of hippocampal silent engrams or the crosstalk between two similar memories through online manipulations, i.e., as mice were engaged in active behavior^14,15,24,54^. This design confines manipulation within a specified temporal window but confounds the effects of engram manipulation with contextual cues. Indeed, in our NOPR paradigm, we successfully maintained the engram in the latent state at recent and remote time points with single and repeated behavioral reminders, respectively (Extended Data Fig.5a, b), but failed to achieve the same result with one week of daily chemo-genetic stimulation (Extended Data Fig.5c-f). Coupled with our successful online rescue (Fig.1a, b), these results suggest that engram manipulation is qualitatively different from behavioral reminders even when the same engram and epoch are used. We thus limited our engram manipulation to at least two hours before mice were introduced to the modified event.

We first assessed whether the enhanced consolidation imparted on the modified event by the silent engram could be weakened if we activate the engram during this phase. We labelled the original NOPR engram with a stimulatory chemo-genetic actuator, then injected CNO or VEH repeatedly on the third and fourth days, until two hours before the modified event (Fig.3a, b). Mice in the VEH group successfully preferred the novel object location in the modified memory test, but the CNO group, which received repeated engram stimulations, did not (Fig.3c and Extended Data Fig.6a, b). Only mice in the CNO group, however, recovered their original memories (Fig.3c and Extended Data Fig.6c, d). This experiment demonstrates that repeated stimulation of the NOPR engram during its silent phase weakened its preferential effect on the consolidation of the modified event.

**Figure 3:**
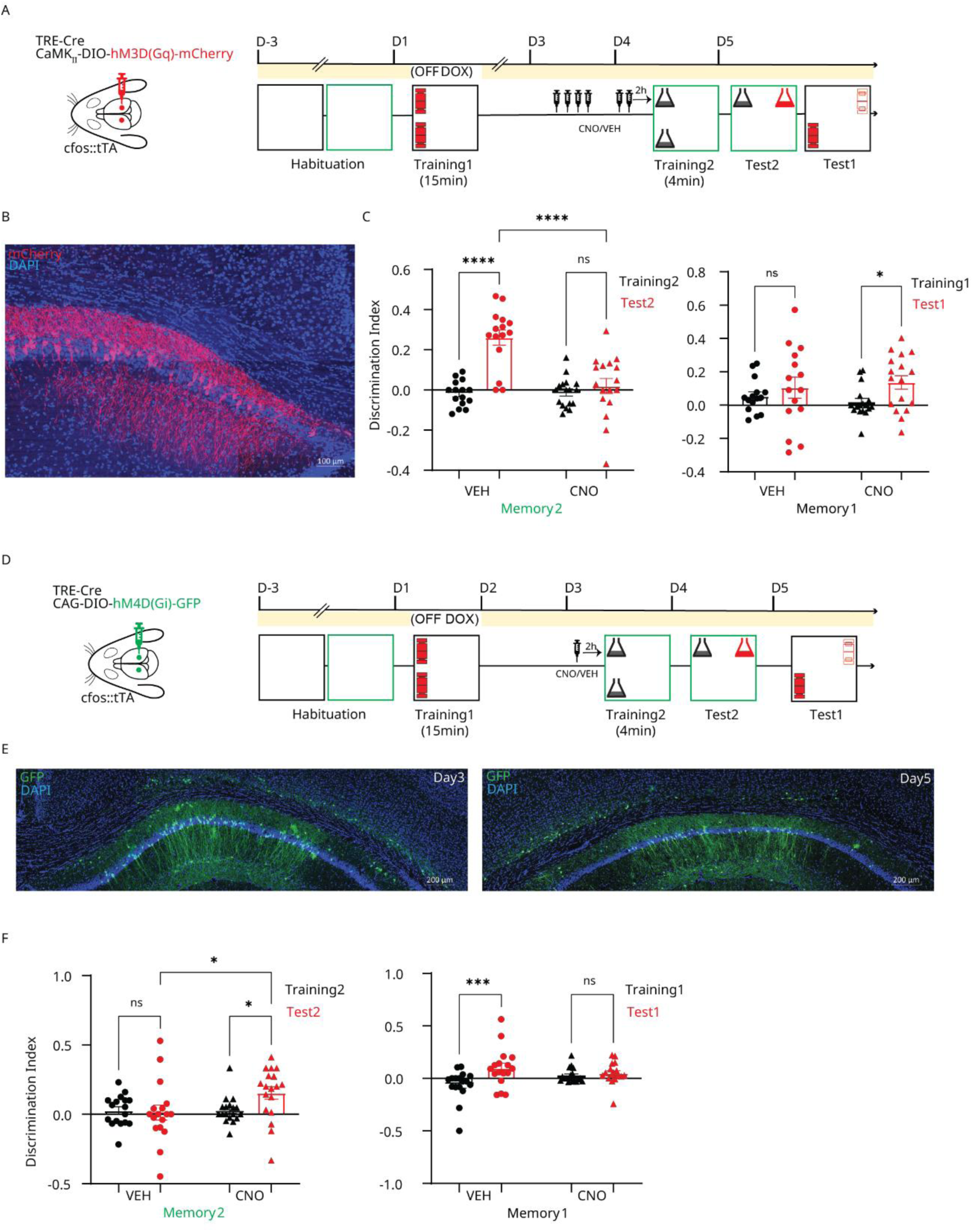
Bidirectional manipulation of NOPR engram. **(A)** Design and behavioral timeline for the chemo-genetic activation of the silent NOPR engram. **(B)** Labelling of engram cells in CA1 with the chemo-genetic activator. Brain sections were immunostained for mCherry (red) and DAPI (blue). Scale bars, 100 μm. **(C)** Discrimination of the novel object’s location at the modified (left) and original (right) memory test for the VEH and CNO groups. *n* = 15 and 17 mice for VEH and CNO, respectively. **(D)** Design and behavioral timeline for the chemo-genetic inhibition of the latent NOPR engram. **(E)** Labelling of engram cells in CA1 with the chemo-genetic inhibitor at Day 3 (left) and 5 (right). Brain sections were immunostained for GFP (green) and DAPI (blue). Scale bars, 200 μm. **(F)** Discrimination of the novel object’s location at the modified (left) and original (right) memory test for the VEH and CNO groups. *n* =17 and 18 mice for VEH and CNO, respectively. All statistical comparisons were performed using repeated measures two-way ANOVA. CNO, clozapine-N-oxide; VEH, vehicle. ns, non-significant; *, P < 0.05; ***, P < 0.001, ****, P < 0.0001. Data are represented as mean ± standard error of the mean (SEM).

Next, we tested whether the latent engram could gain the consolidatory benefit of the silent state if it was inhibited before the second epoch. Our initial experiment showed that the latent phase is present only for two days after encoding (Fig.1a), which left only one day to tag the engram with the inhibitory chemo-genetic actuator, and another for manipulating the engram and performing the modified training (Fig.3d). We confirmed sufficient virus expression was induced at this early time point (Fig.3e), which was critical as only a single injection of CNO or VEH was administered two hours before the modified event. Mice in the VEH group (with an unmanipulated latent engram) performed poorly in the modified memory test, but their CNO counterparts showed significantly better discrimination (Fig.3f and Extended Data Fig.6e, f). In the original memory test, however, only the VEH group mice showed spontaneous recovery, while mice whose engram was inhibited did not (Fig.3f and Extended Data Fig.6g, h).

Finally, we investigated how synaptic GluA_2_-containing AMPARs affect the consolidatory benefit of the silent engram in our paradigm. Blocking the endocytosis of these receptors was shown to maintain the hippocampal engram in a latent state^20^ and to enhance memory retrievability from silent states^24^. Whether these receptors also modulate an engram’s state-induced interaction with different episodes is unclear. We first confirmed that blocking hippocampal endocytosis of GluA_2_-containing AMPARs using the decoy GluA23Y peptide maintains the latent state of the NOPR memory (Extended Data Fig.7). We then exposed mice to our dual NOPR paradigm while micro-infusing either the GluA23Y peptide or its scrambled control, one day after the original event until two hours before the modified one (Fig.4a and Extended Data Fig.8a). GluA23Y impaired the performance in the modified memory test (Fig.4b and Extended Data Fig.8b, c), but exhibited spontaneous recall of the original memory (Fig.4b and Extended Data Fig.8d, e), while the control group did not. Combined, these experiments demonstrate that both direct and indirect manipulations of the NOPR engram before the modified event are sufficient to invert its effects on this weak episode. Our injection schedules were planned to place the modified event well outside the established window for the online administration of CNO^37,55^ and GluA23Y^24^. However, given the low temporal resolution of these manipulations, we can’t exclude the possibility of any residual effects, but we can project that if present, they are less than with strictly online manipulations.

**Figure 4:**
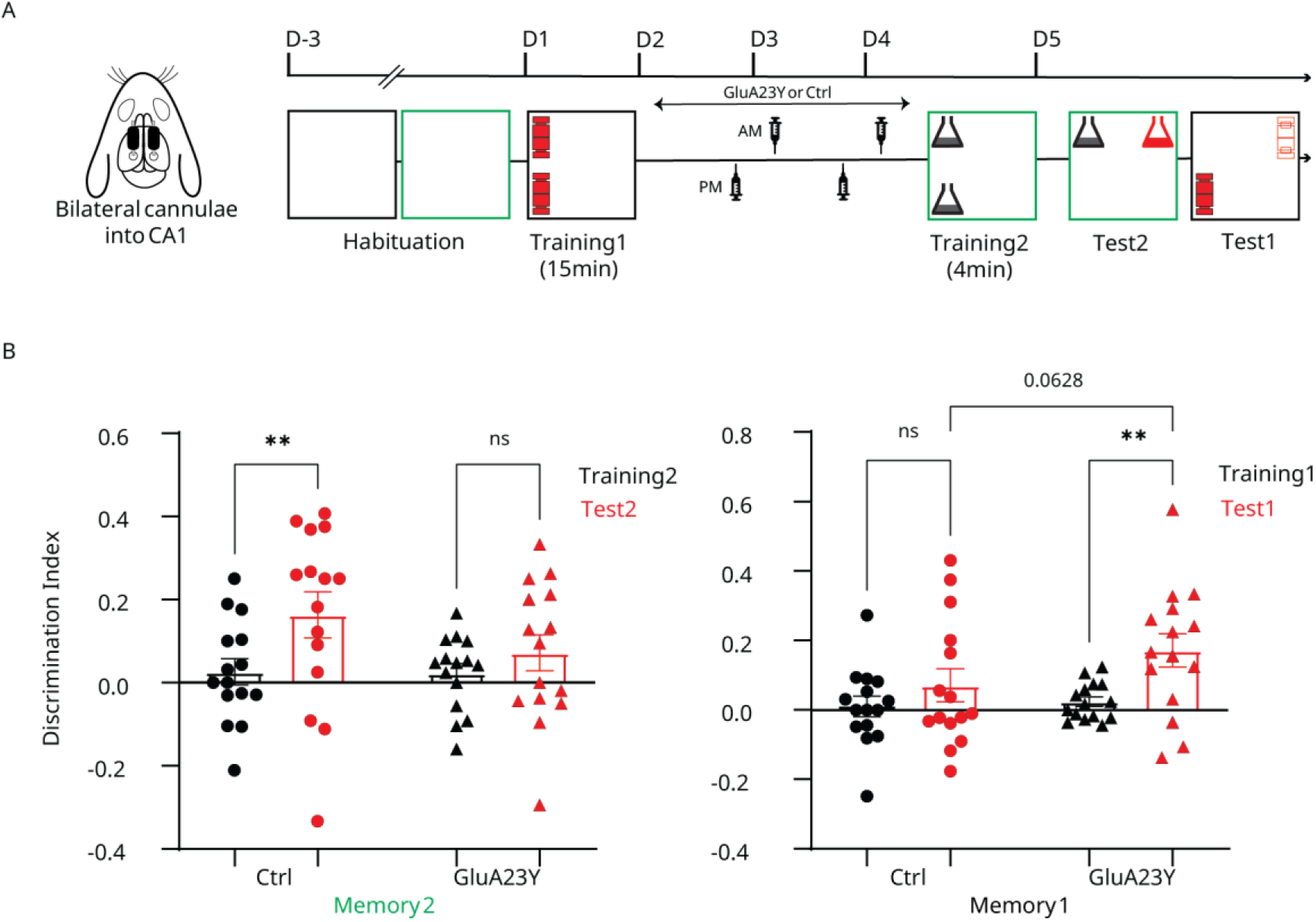
Manipulation of synaptic GluA_2_ AMPARs with the silent NOPR engram. **(A)** Design and behavioral timeline for manipulating synaptic localization of GluA_2_-containing AMPARs in the silent engram protocol. **(B)** Discrimination of the novel object’s location at the modified (left) and original (right) memory test for the Ctrl and GluA23Y groups. *n* = 15 mice per group. All statistical comparisons were performed using repeated measures two-way ANOVA. Ctrl, control Tat-GluA2_3A_ peptide; GluA23Y, Tat-GluA2_3Y_ peptide. ns, non-significant; **, *P* < 0.01. Data are represented as mean ± standard error of the mean (SEM).

### 4. Distinct levels of reactivation for the latent and silent NOPR engrams

We next asked whether latent and silent NOPR engrams displayed distinct reactivation levels when encountering a behavioral event at their respective time points. We first labelled the original NOPR engram with an activity-dependent, color-only (mCherry) tag. Two (latent) or four (silent) days later, we exposed mice to either the modified event or a 4-minute retraining episode, thus having a total of four experimental groups: Latent, Latent Retraining, Silent and Silent Retraining (Fig.5a, b). Mice were sacrificed 90 minutes after the second event to capture its neuronal activity through c-Fos expression^56^. We used a recently developed automated cell counting algorithm^57^ to objectively compare the dynamics of engram reactivation across our four experimental conditions. All groups showed comparable neuronal counts (Fig.5c) and similar, sparse fractions of engram cells (Fig.5d), in agreement with the documented properties of hippocampal memory allocation^2,14,58,59^. When comparing the total fraction of neurons that were activated during the second event (c-Fos^+^ neurons), however, we found that both the Silent and Silent Retraining groups displayed higher activities compared to their latent counterparts (Fig.5e). To understand the relation of this higher activity to the first event, we subdivided all c-Fos^+^ neurons into either non-engram (mCherry^-^) or engram (mCherry^+^) neurons, and rechecked their distribution across our groups. The activation of non-engram neurons was similar between corresponding pairs of latent and silent groups (Fig.5f and Extended Data Fig.9a), indicating that the naïve (non-engram) response to the second event was not affected by the state of the concurrent engram, and that non-engram activity is probably not the source of the observed differences in c-Fos^+^ neurons. Instead, these differences arose mainly from engram (mCherry^+^) neurons, as both silent groups displayed higher c-Fos signal, i.e., engram reactivation, than their latent variations (Fig.5g and Extended Data Fig.9b). All groups showed significant reactivation compared to chance, demonstrating a powerful link between both episodes. Interestingly, all parameters (c-Fos^+^ fraction, non-engram activation, and engram reactivation) were comparable when comparing each engram state to its retraining variation (Extended Data Fig.9c-h), further supporting our conclusion that the observed changes reflect the state of the original NOPR engram. Manual counting also confirmed that the hippocampus displays higher levels of c-Fos when harboring a silent state, that the silent engram is more reactivated than the latent counterpart, and that these changes are state-specific (Extended Data Fig.10). Altogether, this experiment provides objective and quantitative evidence that, when an animal revisits the same event or encounters a highly similar iteration, silent hippocampal engrams are more preferentially recruited than their latent predecessors.

**Figure 5:**
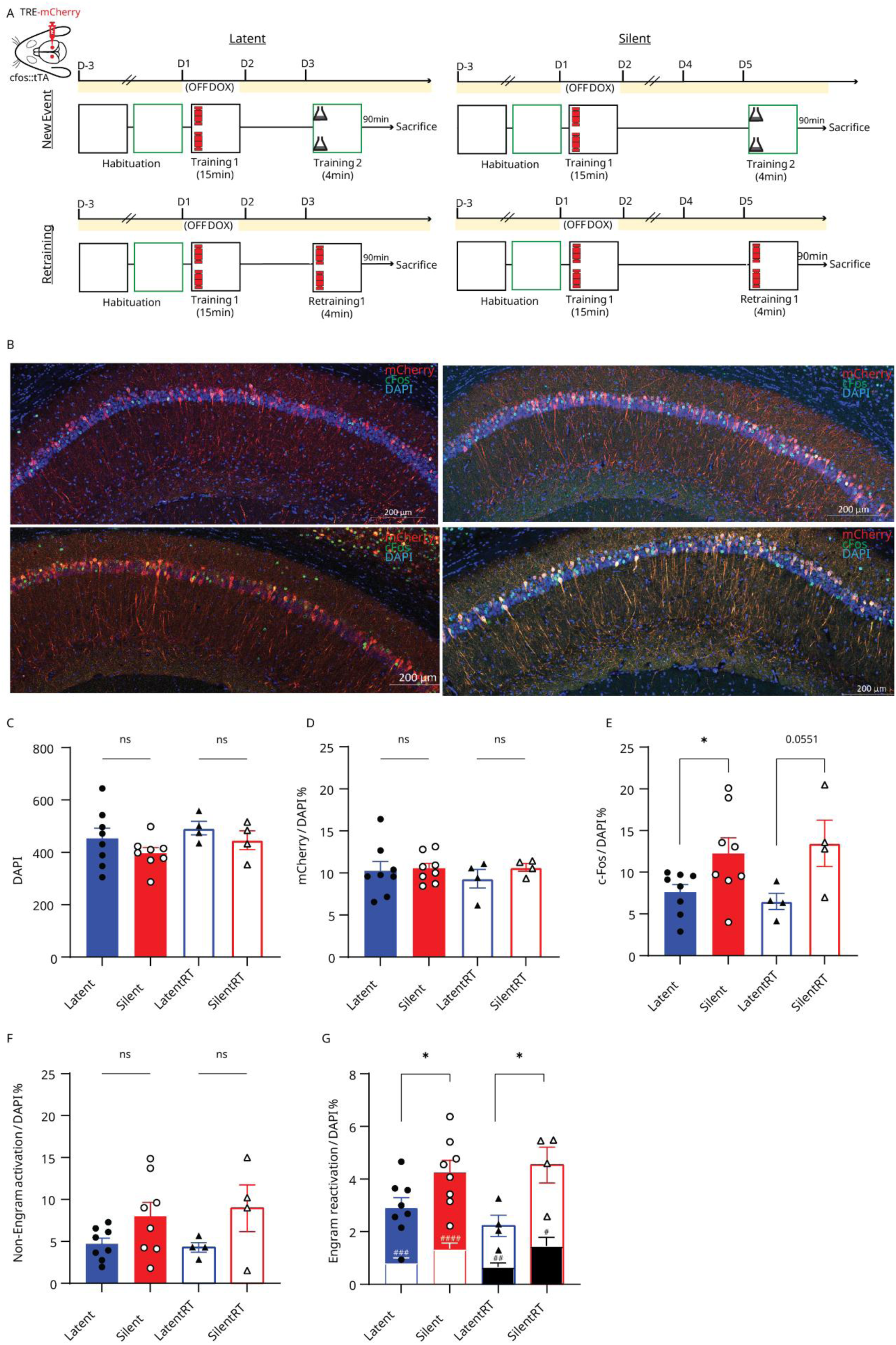
Reactivation of latent and silent NOPR engrams. **(A)** Design and behavioral timeline for the engram reactivation experiment. Latent (left half) and silent (right half) engrams were exposed to the modified event (top half) or a short retraining (bottom half). **(B)** Labelling of engram and active cells in CA1. Brain sections were immunostained for mCherry (red), c-Fos (green) and DAPI (blue). Scale bars, 200 μm. Representative sections are ordered in the same group arrangement as in **(A)**. **(C)** DAPI-positive cells across groups. **(D)** Percentage of engram (mCherry-positive) cells from total counted cells across groups. **(E)** Percentage of active (c-Fos-positive) cells among the total counted cells across groups. **(F)** Percentage of activated non-engram (mCherry-negative, c-Fos-positive) cells among the total counted cells across groups. **(G)** Percentage of reactivated engram (mCherry and c-Fos-positive) cells among the total counted cells across groups. Inner columns represent the chance level for each group. *n* = 8 and 4 mice per group for the modified and retraining events, respectively. All statistical comparisons across groups were performed using an unpaired Student’s t-test, while within-group comparisons against chance level were calculated with a paired Student’s t-test. w. ns, non-significant; *, P < 0.05 for group comparisons and #, P < 0.05; ##, P < 0.01; ###, P < 0.001; ####, P < 0.0001 for comparison to chance level. Data are represented as mean ± standard error of the mean (SEM).

## Discussion

We investigated how silent hippocampal engrams behave when exposed to a highly similar event that can’t produce strong long-term memories on its own. Animals better consolidated this experience when their original engrams were in the silent, not the latent, phase. This outcome depended on the state and was gained and lost when we inhibited and activated the latent and silent states, respectively. Silent engrams were also more strongly reactivated when mice were exposed to a modified event or an exact reminder of the original episode. These results suggest that unique roles and activity dynamics are imparted on hippocampal engrams as they transition from their latent to the silent states. A plethora of clinical^30–35,60,61^ and experimental^12–15,17,19,24,29,54,62^ evidence has shown that the persistence of a memory trace is not always a prelude to its voluntary expression. Silent hippocampal engrams were initially identified by interfering with their consolidation^14,15^, and later proven to be a natural state in systems consolidation, where a mature cortical engram supports memory recall instead of the hippocampal trace^12^. More recently, hippocampal fear memory engrams were shown to exhibit promiscuous activity^10^ during remote recall (and generalization) of conditioned fear memories. These findings indicate that, even with systems consolidation, hippocampal engrams maintain significant activity dynamics, which may reflect an ongoing hippocampal-cortical dialogue to expand, but not dictate, memory expression. Situations when spontaneous recall is not supported by any engram, however, are very different at the experimental level. Both natural forgetting^17^ and memory interference^54^ revealed that reduced hippocampal reengagement is responsible for poor recall, and that it can be modulated to rescue weakened memories.

One major question is whether silent hippocampal engrams still serve a purpose when recall is absent, or do they only serve as an interim state before trace deletion. The persistence of this engram state for several days^14,15^ favors the suggestion that they may still be serving some cognitive purpose(s). In healthy individuals, years-^60,61^ and even a decade^33^-old memories spontaneously re-emerge with similar cues and activities. The hippocampus also plays a significant role in knowledge integration and generalization across experiences^38–46,63–70^. We thus hypothesized that silent hippocampal engrams can serve as a malleable representation to be reused with similar or highly connected events, in a way that is more lenient on the specifics but more widely applicable^36^.

To our knowledge, our study is the first to test for an enduring proactive effect for hippocampal silent engrams on novel events, as previous work mainly focused on the original memory and the means for its recovery^17,54^. Our experimental design provides insight into our results and the possible mechanism(s) at play. For example, a recent study demonstrated that when mice are exposed to two contexts, each with its own set of objects, retroactive interference impairs the original memory by causing intrusions from the second event^54^. This interference is mediated by inter-engram competition, as each event produces its own population, which leads to behavioral forgetting. We believe such competition and interference did not occur in our paradigm for three main reasons. First, we repeatedly habituated mice to both contexts within a short temporal window that induces prospective linking^37,47,50^, whereas the aforementioned study separated contextual learning by 24 hours, which may promote pattern separation. Indeed, our reactivation experiment suggests a strong link at the contextual level, as both the latent and silent engrams were similarly reactivated during both the modified event and the retraining, despite using different contexts and objects in each episode. Another key design difference in our study is that both events are congruent, i.e., simply remembering one event should not impair the performance in the other, as object locations were the same. Lastly, two key results of our reactivation experiment disagree with what we should expect with retroactive interference. Firstly, both the retraining and the new events similarly reactivated the latent and silent engram states in our experiment, while with retroactive interference, re-exposure results in higher engram reactivation than memory update. Secondly, competition between distinct memory engrams occurs with retroactive interference, whereas in our paradigm, non-engram neurons (the pool for a potential competing engram) behaved similarly during both retraining and the modified event. As such, our behavioral design and reactivation results clearly diverge from the expected outcomes of retroactive interference or inter-engram competition.

Another possibility we can debunk in our paradigm is that the strong memory in the latent state proactively impaired efficient encoding of the modified event. Indeed, computational models^71^ as well as human behavioral^72^ and functional magnetic resonance imaging data^73,74^ show that encoding and retrieval act as competing mechanisms in the hippocampus, so that the retrieval of old memories (regardless of its success) impairs the encoding of concurrently experienced information. As we demonstrated in our short-term memory experiment, the state of the hippocampal engram had little effect on learning the second event. Moreover, silent engrams were more strongly reactivated than their latent counterparts, which, if taken as a measure of memory recall, should have further weakened the encoding of the new memory, which we did not observe. Taken together, our behavioral paradigm is unlikely to produce significant memory interference in either direction.

Further dissecting the results of our reactivation experiment may offer better mechanistic insights into how the latent and silent engrams process the modified event. Both states showed significant reactivation (compared to their respective chance levels) with the modified event and retraining epochs, suggesting that this engram response was either mediated by the associated contexts, the spatial object layout, or both. The strength of object memory itself, which is expected to be weaker in the silent state, may not be a significant bottleneck for engram reactivation, as the hippocampus was shown to properly track the quality of contextual information even when the associated item memory is weak^75^, and to develop firing specificity for distinct task structures, rather than absolute environments^76^. Moreover, computational and statistical models support the notion that memories are parsed into a series of latent causes, against which new events are compared to decide whether they fit an already learned structure^77,78^. As such, in addition to the association at the contextual level, both engram states may have correctly referred the second event to the same structure as the original memory, leading to equal reactivation with both the modified event and the retraining episodes. Regardless of the underlying computations, our results show that the state of the hippocampal engram is, by itself, neither a limitation nor a determinant of its ability to compare between epochs.

If both engram states successfully respond to the underlying link between episodes (the contexts, spatial layout, or both), and little to no interference is expected, how can we explain the difference in their behavioral outcomes? Two peculiar findings of our reactivation experiment, namely, higher overall levels of c-Fos and stronger reactivation with silent engrams, may be critically involved. One possibility is that the silent state produces a stronger retrieval of the original memory, as reminders are indeed more effective when memories are weak^79^. A model of the medial temporal cortical regions showed that when stronger memories are recalled, a smaller subset of neurons is more strongly activated than weaker memories, which recruit a broader pool^80^. In our experiment, however, the silent engram displayed no preferential recovery of the original memory compared to the latent state, which should be expected if stronger recall was provoked. Clinical evidence also showed that the degree of original memory reactivation did not differ when subjects succeeded or failed to integrate said memory with a new event^81^, which further supports our claim that retrieval dynamics may be present, but are probably not the key player in our results. The higher reactivation in the silent state may instead reflect a more permissive generalization and integration signal. Clinically, retrieval of older items favors the integration of subsequent ones^82^. Moreover, a hippocampal network model showed that the monosynaptic pathway, when embedded with higher unit activity and neuronal overlap, quickly learns and represents regularities across episodes^46^. Higher hippocampal engram activity is also linked to contextual fear memory generalization during systems consolidation^10^. Hippocampal engrams also contain functionally distinct ensembles, with the c-Fos population promoting memory generalization^55^. Together with these studies, our results support a framework where hippocampal engrams may become more vigilant to a wider array of experiences as they transition into the silent state. This may reflect a distinct purpose for each engram state: latent engrams are primarily concerned with retrieving the original memory, whereas silent, or more remote, engrams primarily aim to integrate and generalize regularities across episodes. Such a graded outcome for engram (re)activation was previously reported^15^, and echoes our model of memory linking, where the degree of engram overlap between two episodes, rather than its existence *per se*, determines the cognitive outcome^50^. As we discussed earlier, it is unlikely that our results stem from inter-engram competition^83^, but within-engram communication is a novel and promising candidate. Indeed, the rigidity of hippocampal encoding can be relaxed by manipulating neurons in the superficial layer^63^, which are also well positioned to exert feed-forward inhibition on the deep layer^84^, and inhibitory transmission can alter the degree of engram selectivity^85^.

Our study reveals a previously overlooked dynamic of hippocampal memory processing, where silent engrams remain responsive to contextually and semantically linked events. Whether distinct neuronal populations drive the responses of latent and silent engram states remains an exciting avenue for future research. Testing the cognitive reach of these vigilant representations, both within and outside the hippocampus, will also extend our understanding of how higher-order memory integration operates across the countless experiences we accumulate throughout life.

## Methods

### Animals

Male C57BL/6J mice (Sankyo Labo Service Co. Inc., Tokyo, Japan) were used throughout this study, except for the engram manipulation experiments, for which we used cFos-tTA transgenic mice, where the expression of tetracycline-controlled trans activator (tTA) is controlled by the activity-dependent promoter of the immediate early gene *c-fos* (Mutant Mouse Regional Resource Centre, stock number: 031756-MU)^86^. Transgenic mice were maintained on 40 parts-per-million doxycycline (DOX) pellets until 48 hours before the engram labelling session, at which point they were switched to regular mouse chow. Four to six hours after labelling, mice were fed 1000 parts-per-million DOX pellets until sacrifice. Mice used for behavioral analyses were 12–20 weeks old, maintained on a 12 h light/dark cycle at 24°C ± 3°C and 55% ± 5% humidity, and had access to food and water *ad libitum*. Following surgeries and before any behavioral procedures, mice were individually housed for a week. All procedures involving the use of experimental animals obeyed the guidelines of the National Institutes of Health and were approved by the Animal Care and Use Committee of the University of Toyama.

### Viral construct

Plasmids for the AAV9-TRE3G-Cre were obtained as described previously^86^. The plasmids for AAV9-hSyn-DIO-hM3d(Gq)-mCherry and rAAV-CAG-FLEX-rev-hM4D(Gi)-2a-GFP were obtained from Addgene (plasmid #44361 and #52536, respectively). Plasmids for the AAV9-CaMKII-DIO-hM3d(Gq)-mCherry were prepared from those of AAV9-hSyn-DIO-hM3d(Gq)-mCherry and pAAV-CaMKII-hM4D(Gi)-mCherry from Addgene (plasmid #50477). AAV vectors were produced using a previously described protocol^87^. AAV9-TRE3G-Cre was prepared at 2.35E+12 copies/µl and was used at a 1400x dilution throughout this study. AAV9-hSyn-DIO-hM3d(Gq)-mCherry was prepared at 4.55E+11 copies/µl and diluted ten-fold before injection. The remaining viruses, namely AAV9-CaMKII-DIO-hM3d(Gq)-mCherry and rAAV-CAG-FLEX-rev-hM4D(Gi)-2a-GFP, were prepared at 5.77E+10 and 6.32E+11 copies/µl, respectively, and used without dilution. For the engram reactivation experiment, AAV9-TRE3G-mCherry was used without dilution at 4.74E+14 pfu/ml.

### Drugs and peptides

For the DREADD experiments, Clozapine -N-Oxide (CNO, Sigma Aldrich, Japan Co., Tokyo, Japan) was dissolved in dimethyl sulfoxide to prepare a 10x stock solution, and frozen at −30°C. Before use, 1 ml of the stock solution was diluted with 9 ml of saline to prepare a final solution of 10% dimethyl sulfoxide in saline, which was intraperitoneally (i.p.) injected at a dose of 0.5 mg Kg^-1^ and 4 mg Kg^-1^ for the excitatory and inhibitory DREADDs, respectively^37,55^. For the synaptic GluA2s experiments, both GluA2_3Y_ (L-amino acid sequence YGRKKRRQRRRYKEGYNVYG) and the CTRL peptide (L-amino acid sequence YGRKKRRQRRRAKEGANVAG) were obtained from AnaSpec (Funakoshi Co., Ltd., Tokyo, Japan) and prepared as described previously^88^. Briefly, peptides were dissolved in phosphate-buffer saline into single-experiment volumes and stored at −80°C. Immediately before use, aliquots were diluted to a final concentration of 40 ng µl^-1^, and 1 µl was injected into each hippocampus per dose. After the last behavioral session, mice were anaesthetized, and their hippocampi were bilaterally injected with rhodamine. Fresh brains were then harvested to confirm cannula placement and verify the spread of rhodamine in both hippocampi. Only mice with correct placement and spread were included in the study.

### Surgery

Mice were 10–14 weeks old at the time of surgery. They were initially anesthetized with 5% isoflurane, then received i.p. injections of a triple mixture containing Medetomidine (0.75 mg Kg-1, Orion Corporate, Finland), Butorphanole (5 mg Kg-1, Meiji Seika Pharma Co., Ltd., Tokyo, Japan), and Midazolam (4 mg Kg-1, Sandoz, Japan Co., Tokyo, Japan). Mice were then placed in a stereotactic apparatus (Narishige, Tokyo, Japan), and a cranial window was drilled above both hippocampi. For the DREADD experiments, 0.6 µl of virus mixture was bilaterally injected into the CA1 region (+2.2 mm anteroposterior [AP], ±1.4 mm mediolateral [ML], +1.5 mm dorsoventral [DV] from Bregma) at a rate of 0.1 µl/s using a micro syringe pump attached to a glass capillary tube (Narishige). Injection started three minutes after inserting the capillary tube, which remained in place for ten minutes following termination of injection to ensure proper virus diffusion. For the GluA_2_-AMPARs experiments, stainless guide cannulas (Plastics One, Roanoke, VA, USA) were placed at +2.2 mm AP, ±1.4 mm ML, +1.0 mm DV from Bregma, and fixed to anchor screws drilled into the skull beforehand using dental cement. Dummy cannulas (Plastics One) were then inserted into the guide cannulas to prevent clogging. Following surgery, all mice received Atipamezol (0.75 mg Kg-1, Orion Corporate, Finland) and were placed on heating pads for two days or until visible recovery. For GluA2_3Y_ and CTRL peptide injections, mice were kept under 1.5% isoflurane anesthesia, dummy cannulas were removed, and a stainless-steel injection cannula (Plastics One) protruding 0.5 mm from the guide cannula was used to inject the peptides at a rate of 0.2 µl/min. following the termination of the injection, cannulas were left for five minutes. Mice that did not fully recover from surgical and injection procedures were excluded.

### The novel object place recognition paradigm

Mice were individually housed one week before the start of all behavioral experiments. The behavioral soundproof room was adjacent to the mice maintenance room, and mice were transported to the behavioral room for 30 min before any procedure. The context used for the novel object place recognition (NOPR) was a grey square (320 mm × 320 mm) with a grey acrylic floor. The walls were covered with off-white tape to reduce reflection and glare. For the original NOPR training, the right and left walls of the context, with respect to distal room cues, were covered with distinct vertical patterns (black lines and circles on a white background), and the objects were red ceramic models of a Japanese post-box (200 mm height × 50 mm width). For the second (modified) NOPR training, the walls and floor were covered with green plastic flooring with a pointy texture. Wall patterns were also changed (horizontal white lines and stars on a green background), and the context was rotated 90° so that patterns were on the front and back walls with respect to distal room cues. Objects for the modified training consisted of standard glass flasks (150 mm height × 80 mm diameter at the base) filled with odorless grey cement. We performed NOPR training and testing as described previously^89^, with a few modifications. Before the start of the experiment, mice were habituated to the experimenter, the behavioral room, and the context for four consecutive days. Each habituation session consisted of six minutes of handling followed by a single ten-minute session of free context exploration without any objects. Mice receiving the two-memory protocol had two context habituation sessions each day, one for each context. Following the last day of habituation, transgenic mice were taken off DOX and kept in the maintenance room for 48 hours, while C57BL/6J mice proceeded directly to the strong training on the next day. For the training sessions, two identical copies of the context-respective object were placed in different corners. Mice were gently held by the base of the tail and introduced to the context, facing a corner that would remain empty during training and testing, to avoid any object preference or avoidance due to handling. Mice were then allowed to explore for fifteen minutes (original training) or four minutes (modified training) before being returned to their home cages. With respect to distal room cues, object locations were similar for each mouse in both training events, but counterbalanced across all mice. Contexts and objects were cleaned with 70% ethanol, followed by water, then dried carefully to remove any olfactory cues between mice. Test sessions proceeded similarly, except that for each mouse and for each memory test, one object was moved to a novel location. The novel location was the same while the familiar location was different in the original and modified memory tests, such that for each mouse and memory, the final test layout was unique. Object locations and training and test layouts were counterbalanced across mice.

### Behavioral analysis

All sessions were captured with an overhead web camera (Logitech HD Pro C920) mounted on a vertical stand that rose one meter above the context. For all mice, scoring was manually performed by the same experimenter, who was blind to the objects’ location identities but not to the mouse group/treatment. A mouse was considered to be exploring an object when it faced the object within a two-centimeter distance or when it directly touched that object. Complete immobility near the object, climbing above the object, or getting stuck between the object and the wall were not considered exploration. We calculated location preferences for training and test sessions as previously described^89^. Briefly, to quantify the fraction of time the mouse spent exploring each object during training, a preference index was calculated as follows: Object preference index = (time spent exploring one object / total time spent exploring both objects). Additionally, a discrimination index was used to quantify how much the mouse preferentially explored one object over another, and was calculated as follows: Object discrimination index = [(time spent exploring that object -time spent exploring the other object)/ (total time spent exploring both objects)]. The preference index was used to confirm that mice exhibited no location bias during training events, with an exclusion cutoff of 65%. The discrimination indices for respective training and testing sessions were compared to evaluate preferential exploration of the novel location at test. Mice that explored for less than 15 seconds during any training or test session were excluded. The object that will change location is annotated as Object 1.

### Immunostaining

Mice were deeply anesthetized using a triple anesthesia mixture and transcardially perfused with phosphate buffer saline (PBS), followed by 4% formaldehyde in PBS (PFA) solution. Brains were carefully extracted from the skull and post-fixed in PFA overnight. The following day, brains were equilibrated in a 25% sucrose in PBS solution for 24 hours before being carefully dried and stored at –80°C until further processing. Floating 50 µm sections were cut using a cryostat and initially allowed to rest in room temperature PBS solution in a 12-well cell culture plate (Corning, NY, USA). Sections were then placed in a blocking solution consisting of 3% normal donkey serum (S30, Chemicon by EMD Millipore, Billerica, MA, USA) in PBS containing 0.05% Tween 20 and 0.2% Triton X-100 (PBST) at room temperature for 1 hour. Sections were then washed three times with PBST before being incubated in a blocking solution with one or more of the following antibodies: rabbit anti-cFos (1:1000, Santa Cruz Biotechnology, sc-52), rat anti-cFos (1:5000, Synaptic Systems, 226017), rabbit anti-mCherry (1:1000, Clontech, 632496), and rat anti-GFP (1:1000, Nacalai Tesque, 04404-84, GF090R). The plate harboring the sections was then wrapped in parafilm and aluminum foil and incubated at 4°C for 36 hours. Following the primary incubation, sections were washed three times with PBST before being incubated in a secondary blocking solution containing DAPI (1 μg/ml, Roche Diagnostics, 10236276001), as well as one or more of the following antibodies: donkey anti-rabbit IgG Alexa Fluor 488 (1:1000, Invitrogen, A21206), donkey anti-rabbit IgG Alexa Fluor 546 (1:1000, Invitrogen, A10040), and donkey anti-rat IgG Alexa Fluor 480 (1:1000, Invitrogen, A21208). Finally, sections were washed three times with PBST, then once with PBS, and mounted on glass slides with Prolong Gold anti-fade reagent (Invitrogen).

### Confocal microscopy

Fluorescent images were acquired using an HS All-in-one Fluorescence Microscope BZ-9000E (Keyence, Illinois, USA) with 10 and 20× plan apochromat objective lenses. Z-stack fluorescent images were acquired using a Zeiss LSM 780/900 confocal microscope (Carl Zeiss, Jena, Germany) with a 10 and 20× plan apochromat objective lens. All acquisition parameters, including photomultiplier tube assignments, pinhole sizes, and contrast values, were kept constant within each magnification.

### Quantification of engram reactivation

Ninety minutes after the last behavioral session, mice were anesthetized and transcardially perfused as described earlier to capture and fix c-Fos signal^90^. For each mouse, fluorescent images were acquired from 5-20 z-stacks per section, and 3-4 sections were captured per mouse.

#### A. Automated counting

We used an automated cell detection algorithm to detect DAPI-positive mCherry and c-Fos signals and their overlap, as previously described^57^, with two main modifications. First, we set all fluorescence levels to the “Best Fit” setting using ZEN 2.5 blue edition software (Zeiss). This ensured that every image was unbiasedly set to an automatically fixed setting before being fed to the detection algorithm. Second, we set the detection threshold for all channels to 10 (instead of the original level of 5). This was to reduce a specific discrepancy we observed with lenient thresholds, where the detection of either mCherry or c-Fos overlap with DAPI sometimes exceeded their maximum counts. Doubling the detection threshold corrected that error across all our experimental groups and was thus used for all analyses.

The chance level for engram reactivation as a fraction of the total population was calculated as described previously^86^, by multiplying the population percentages of mCherry- and c-Fos-positive neurons. When measuring reactivation as a fraction of the engram population, the chance level was corrected to that of c-Fos-positive neurons.

#### B. Manual counting

A rectangular 1000μm area was drawn to include most of the CA1 pyramidal cell layer. Positive signals within this area were then marked and counted separately for mCherry and c-Fos, with the other channel invisible to avoid bias (DAPI was always on to validate signals). After both channels were individually marked and counted, their signals were overlaid, and overlapping cells were marked and verified. For all channels, signals that do not match known cellular morphology or could not be strongly dissociated from surrounding noise were omitted.

### Statistics

All statistical analyses were performed using Prism 8 (GraphPad Software, San Diego, CA, USA). Multiple-group comparisons were made using a two-way repeated measures analysis of variance (ANOVA) with Sidak’s *post hoc* test. Data from the engram reactivation experiment were analyzed with either an unpaired (across groups) or a paired (within group) Student’s *t*-test

**Table 1.**
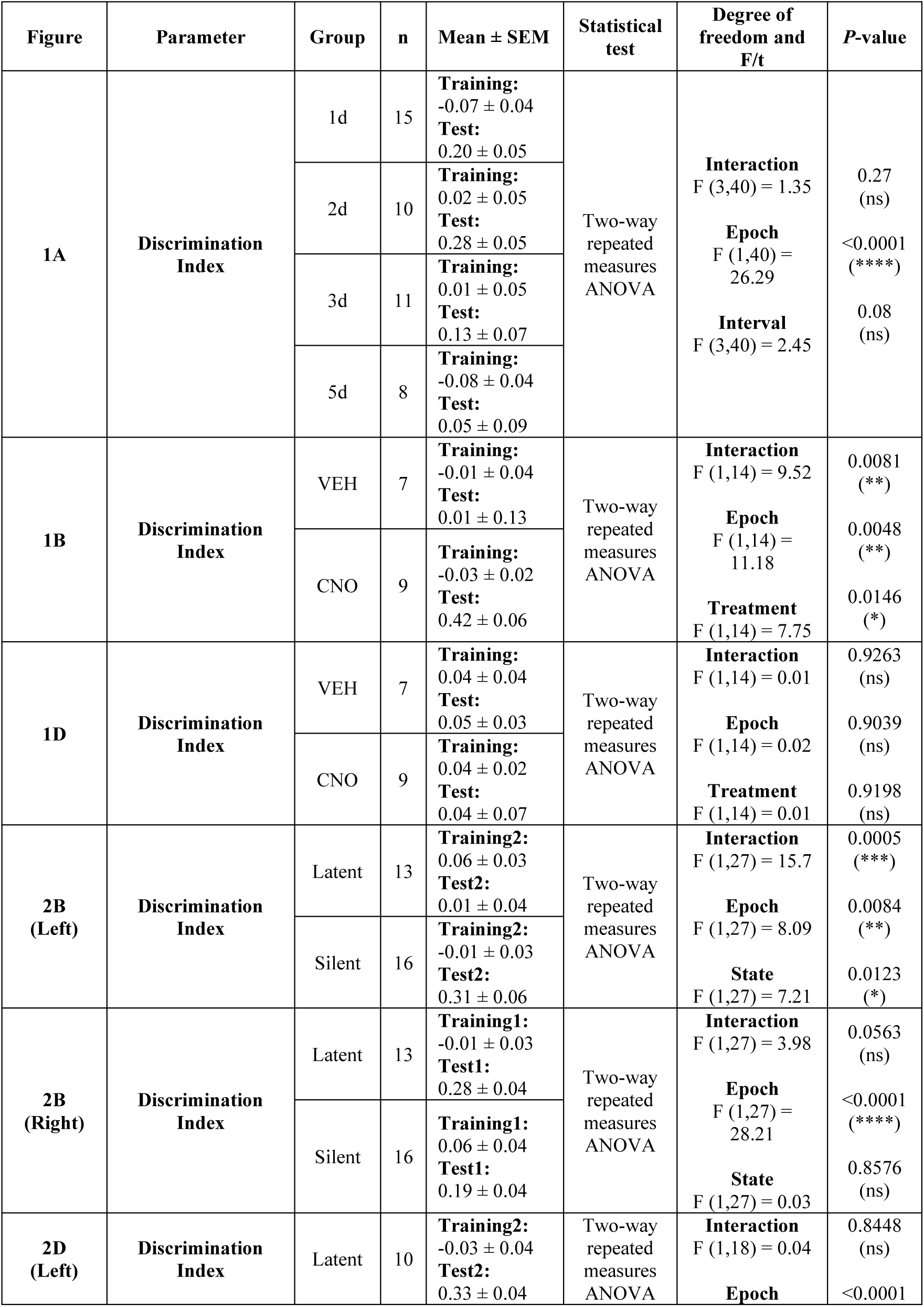
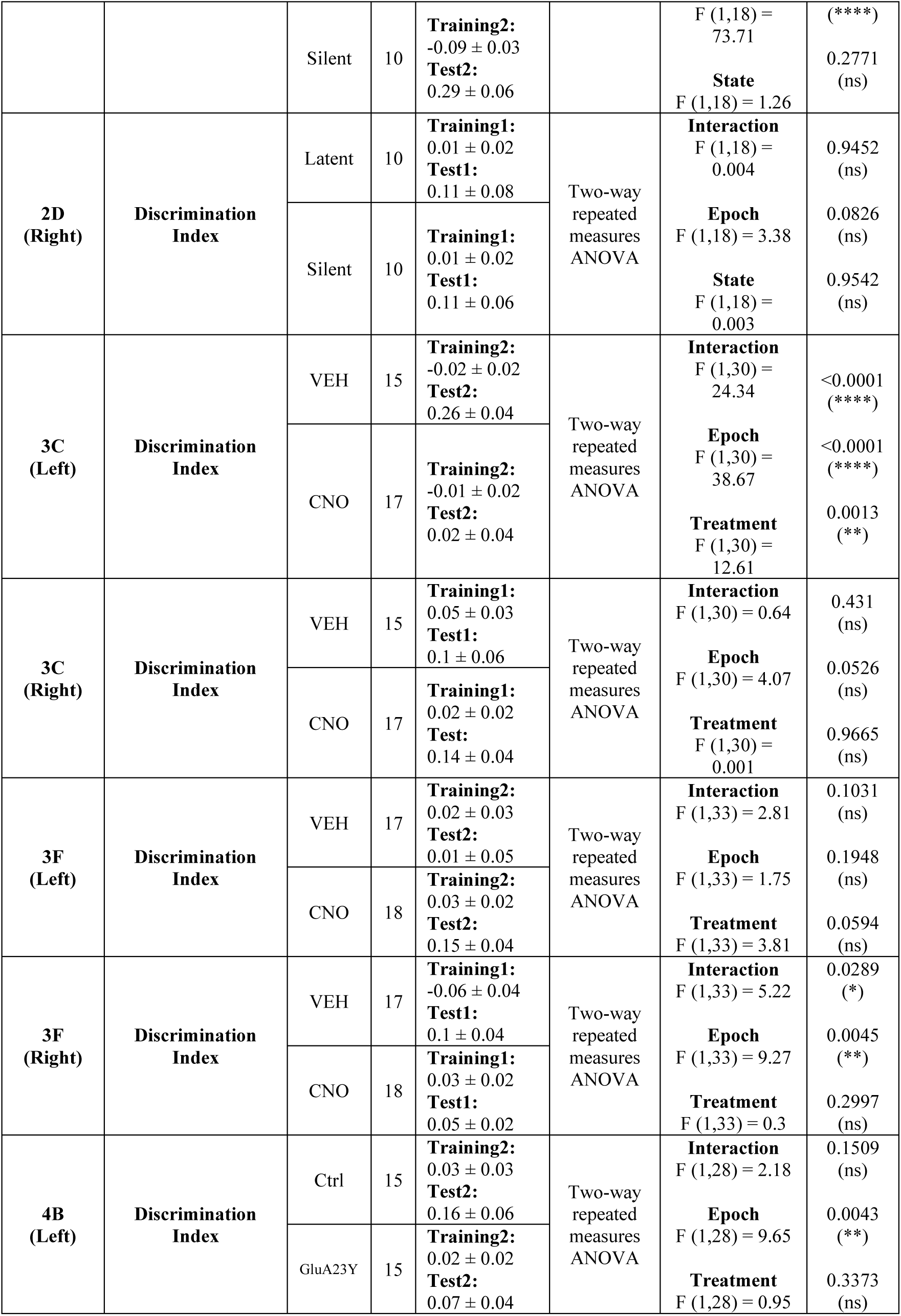
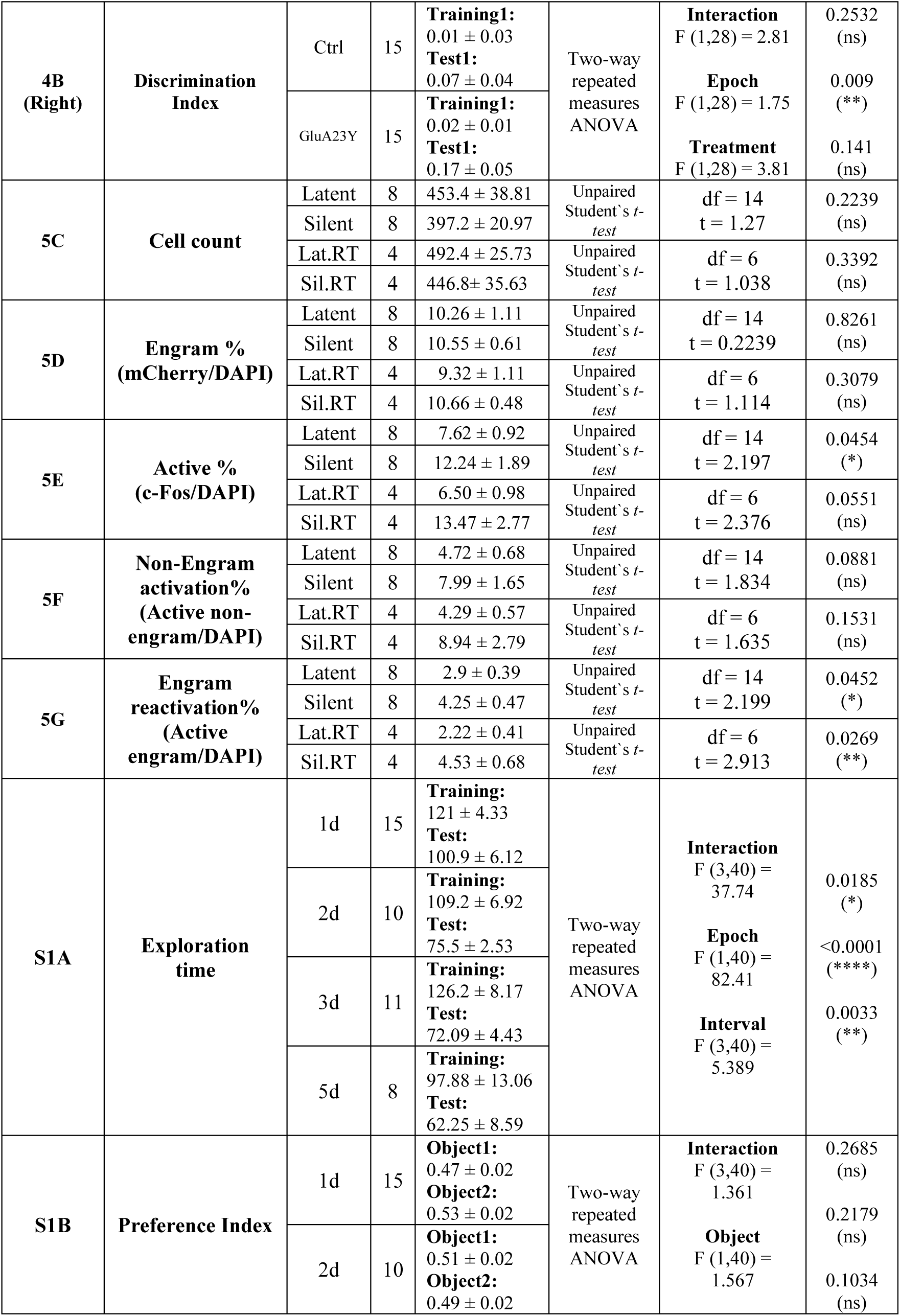
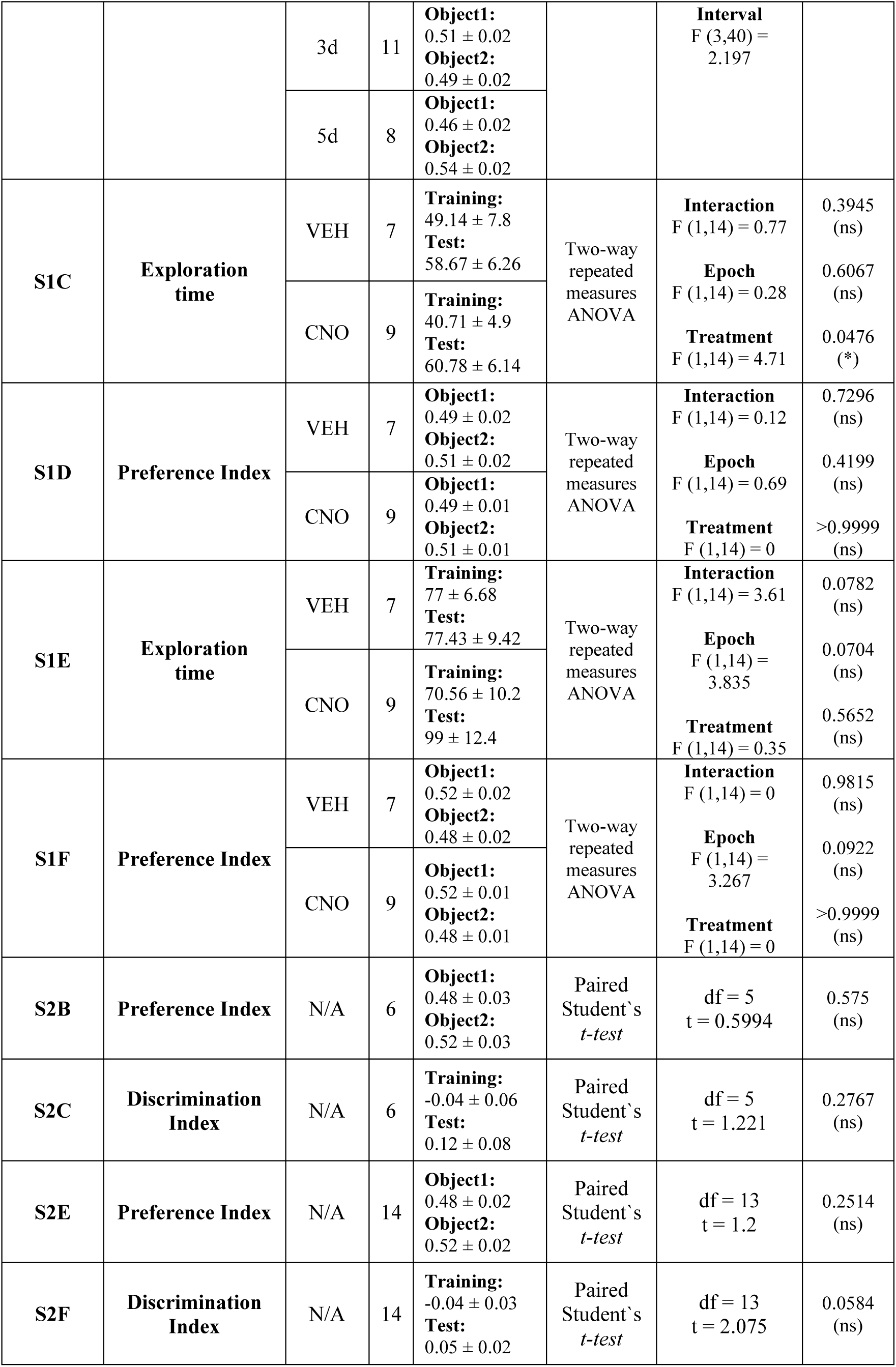
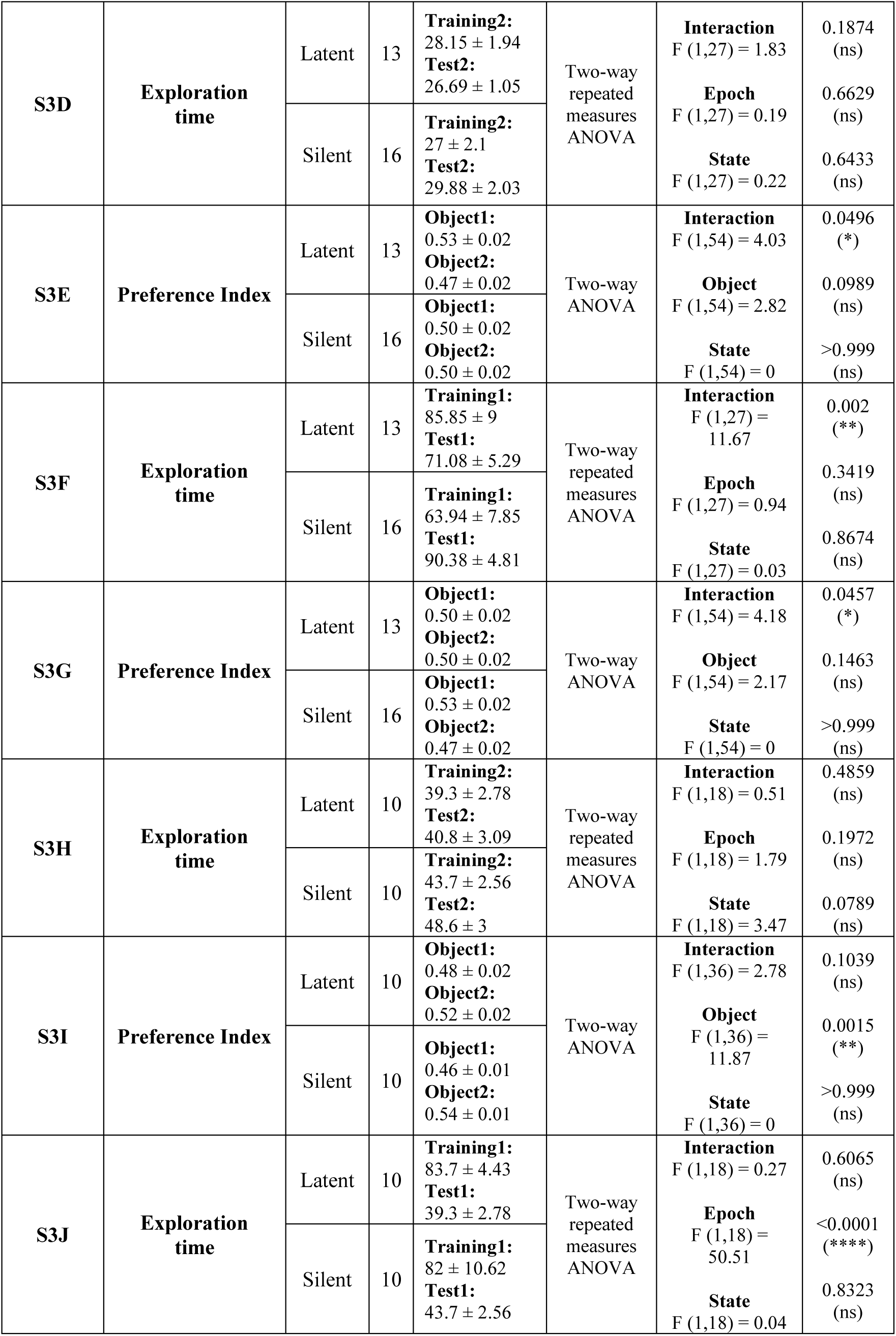
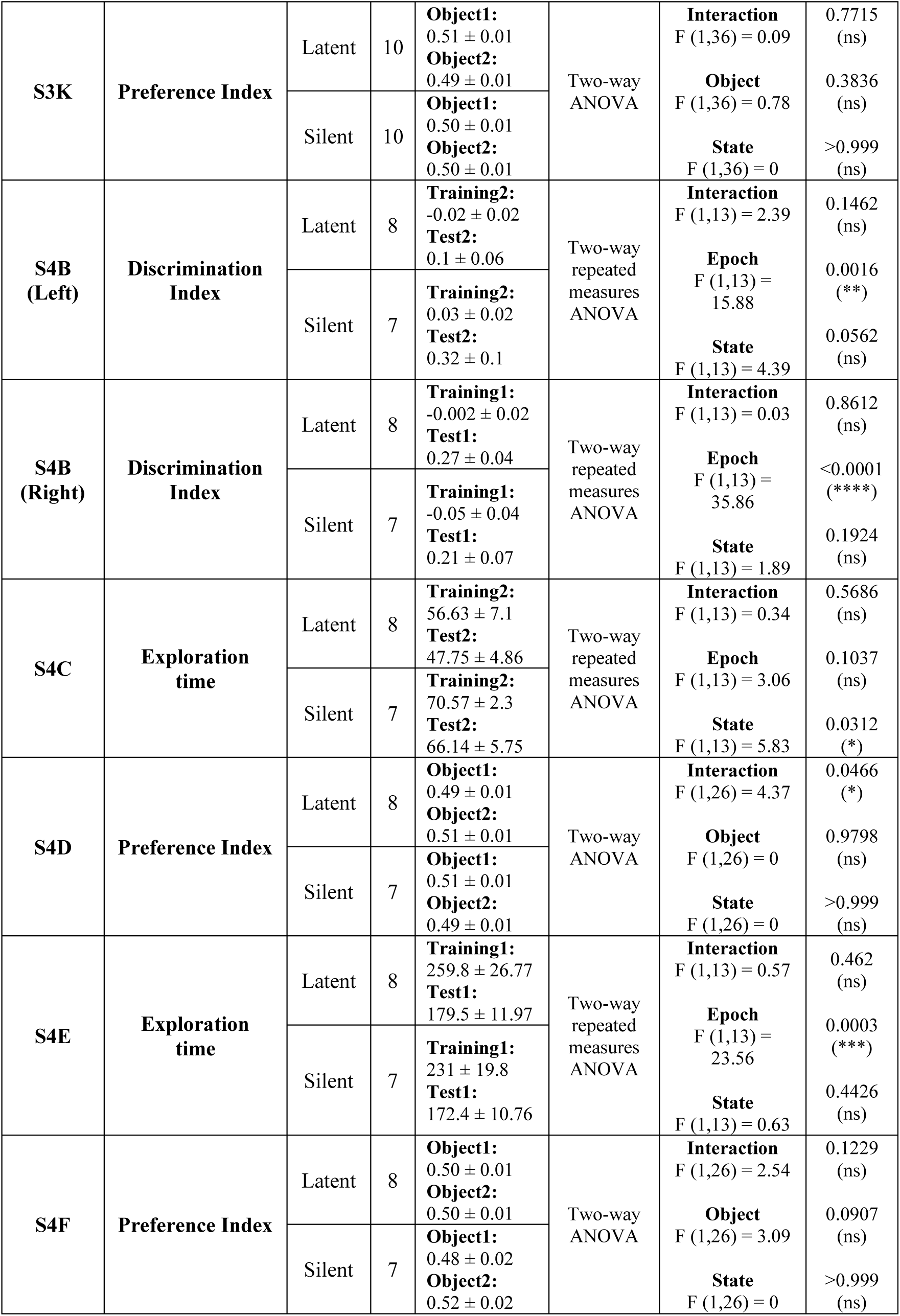
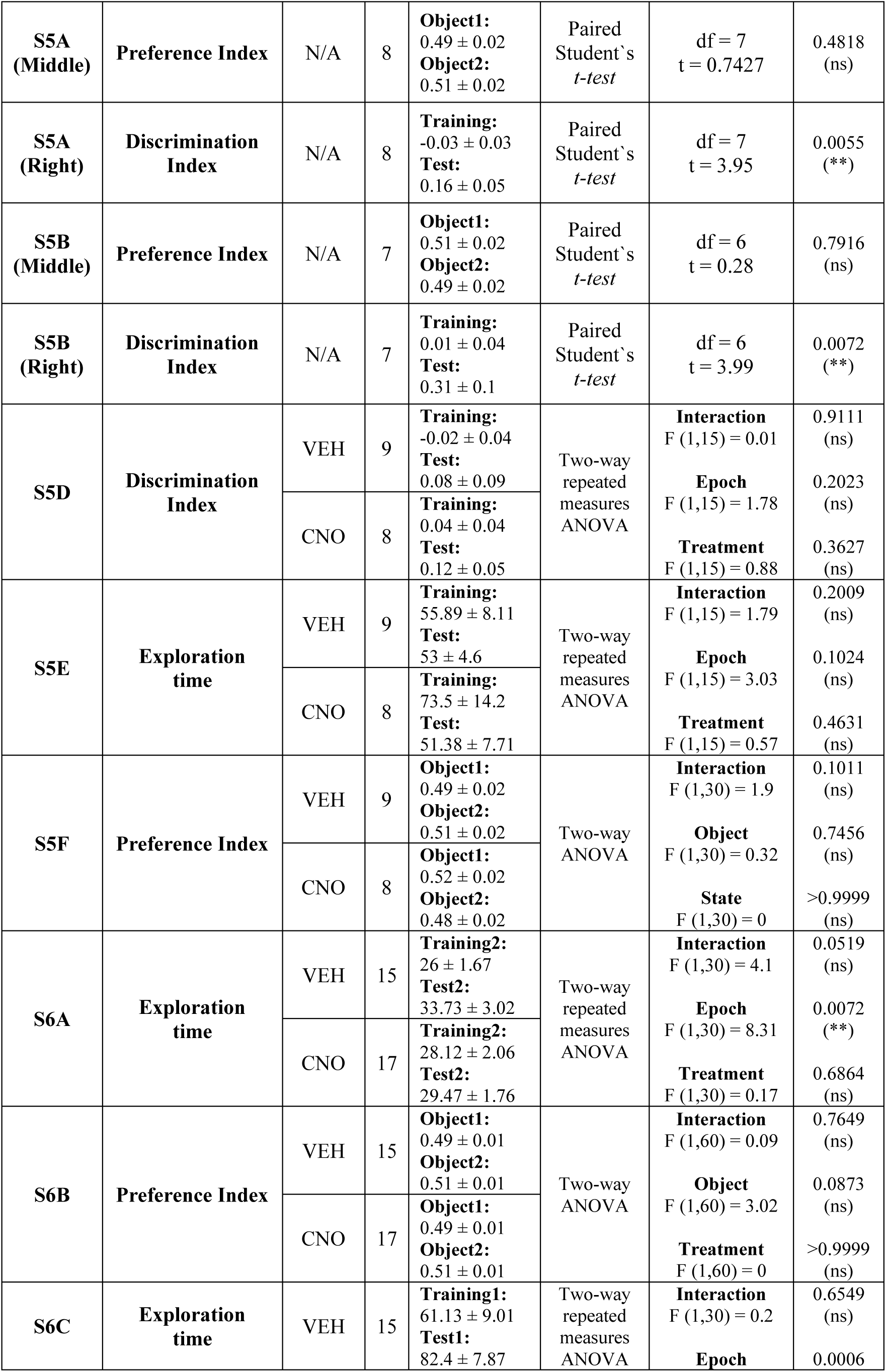
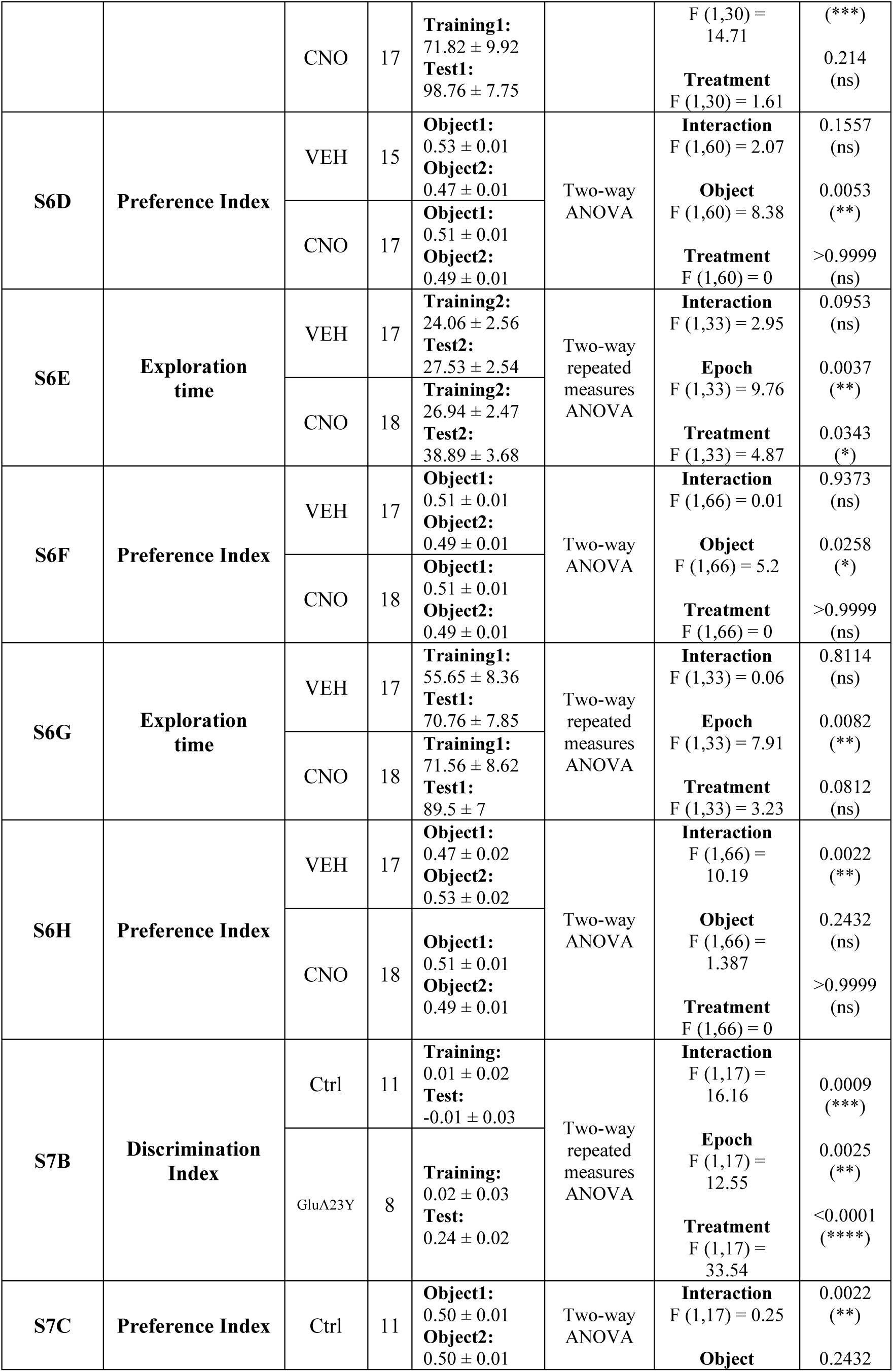
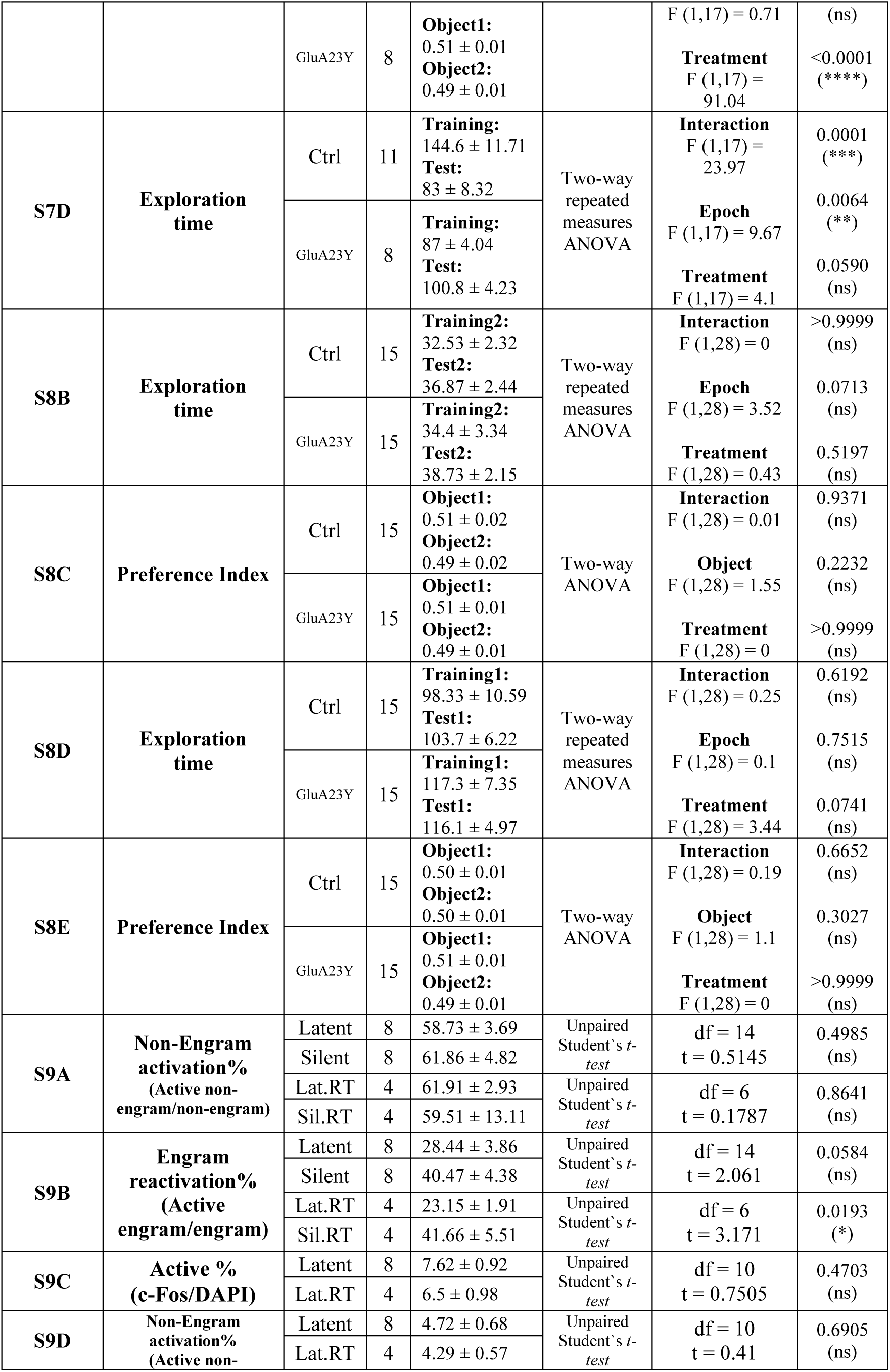
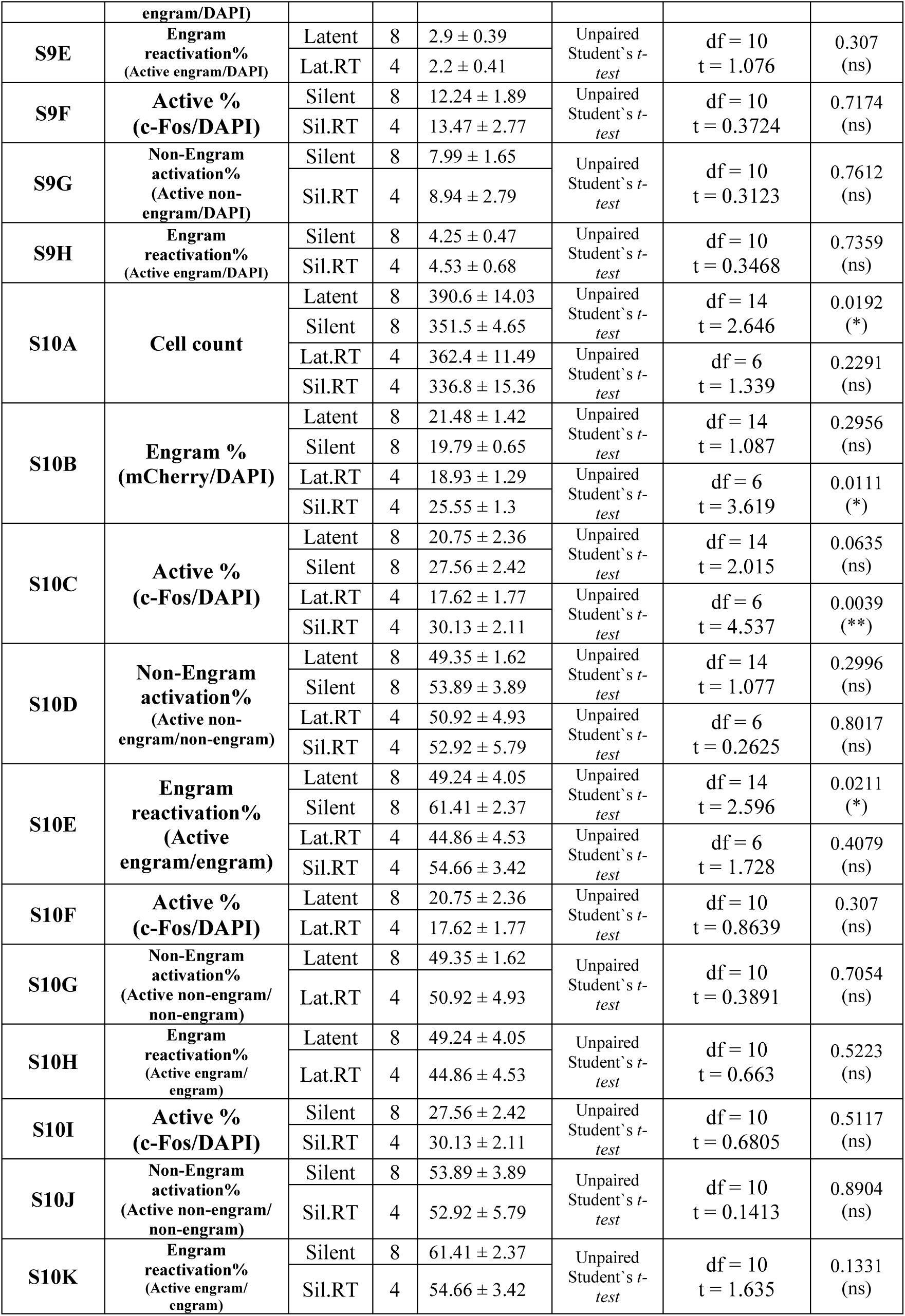
Statistical result summary.

## Acknowledgments

We thank Reiko Okubo-Suzuki and Emi Murayama for constructing the viral vectors used in this study. We also acknowledge Noriaki Ohkawa, Akinobu Suzuki, Jun Yokose, Hirotaka Asai, Md Jahangir Alam, Khaled Ghandour, Mostafa R. Fayed, and Ahmed I. Zaher for their valuable discussions and contributions. We thank all members of the Inokuchi laboratory at the University of Toyama.

## Author Contributions

A.C., K.A., and K.I. conceptualized the study; A.C. performed surgeries, behavioral experiments, immunohistochemistry, and analyses. A.C. and K.I. prepared the manuscript. K.I. supervised the entire project. This work was supported by JSPS KAKENHI (grant number JP18H0521, JP23H054476), the Core Research for Evolutional Science and Technology (CREST) program (JPMJCR13W1 and JPMJCR23N2) of the Japan Science and Technology Agency (JST), a Grant-in-Aid for Scientific Research on Innovative Areas “Memory dynamism” (JP25115002) from MEXT, and the Takeda Science Foundation to K.I. This work was also supported by JSPS KAKENHI (grant number JP19K16892) to K.A. and Uehara Memorial Foundation scholarship to A.C.

## Availability of data and materials

All necessary data is provided within the main manuscript, Extended Data and the source file.

---

**Extended Data Figure1 (Related to Fig.1).**
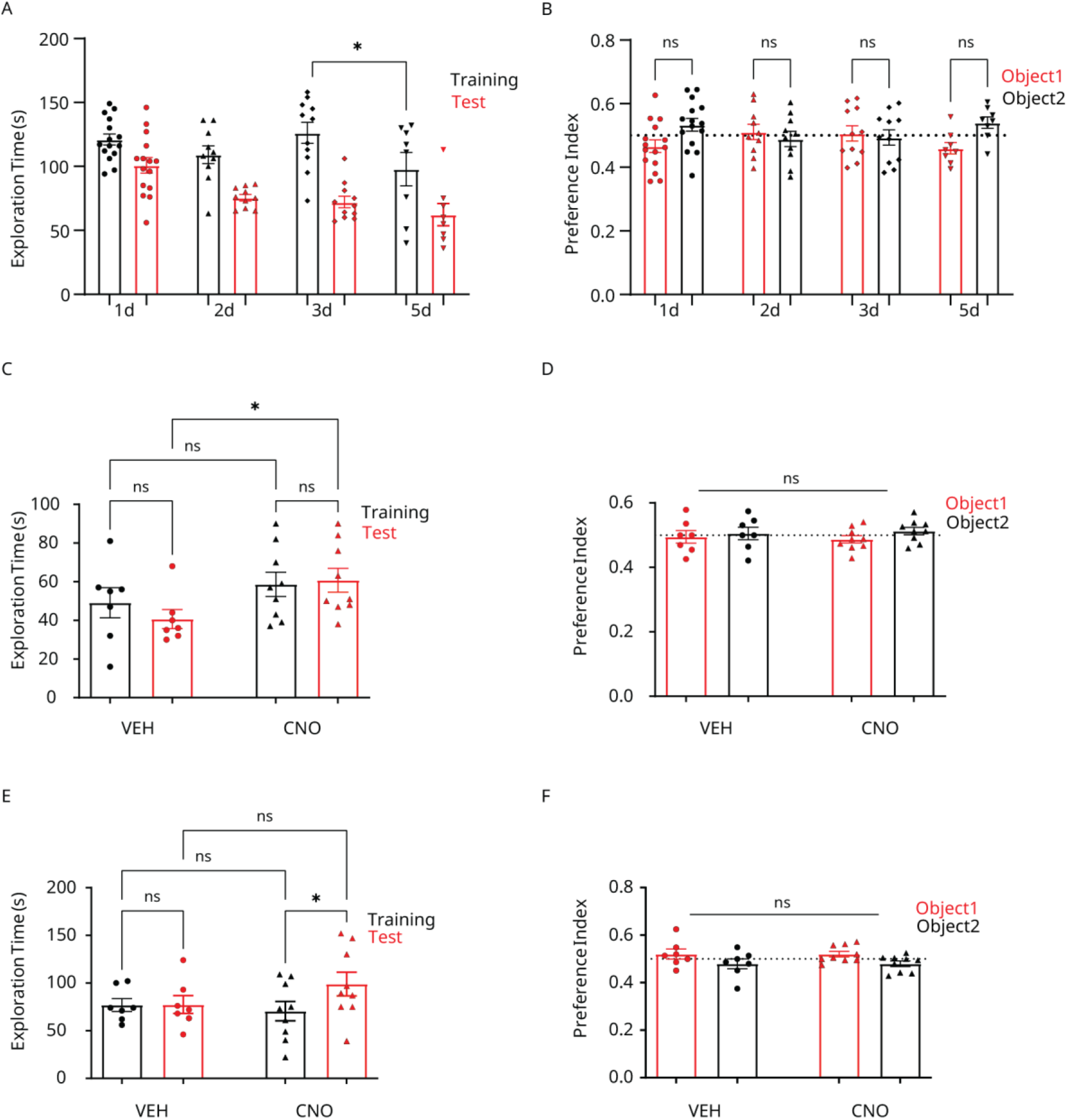
**(A)** Exploration times during training and testing for mice in Fig.1A. **(B)** The preference index for both objects during training. *n* = 15, 10, 11, and 8 mice for 1, 2, 3, and 5 days, respectively. **(C)** Exploration times during training and testing for mice in Fig.1B. **(D)** The preference index for both objects during training for mice in Fig.1B. *n* = 7 and 9 mice for VEH and CNO, respectively. **(E)** and **(F)** Similar to **(C)** and **(D),** but for the chemo-genetic rescue experiment at 28 days for mice in Fig.1D. *n* = 7 and 9 mice for VEH and CNO, respectively. All statistical comparisons were performed using repeated measures two-way ANOVA. CNO, clozapine-N-oxide; VEH, vehicle. ns, non-significant; *, *P* < 0.05. Data are represented as mean ± standard error of the mean (SEM).

**Extended Data Figure2 (Related to Fig.2).**
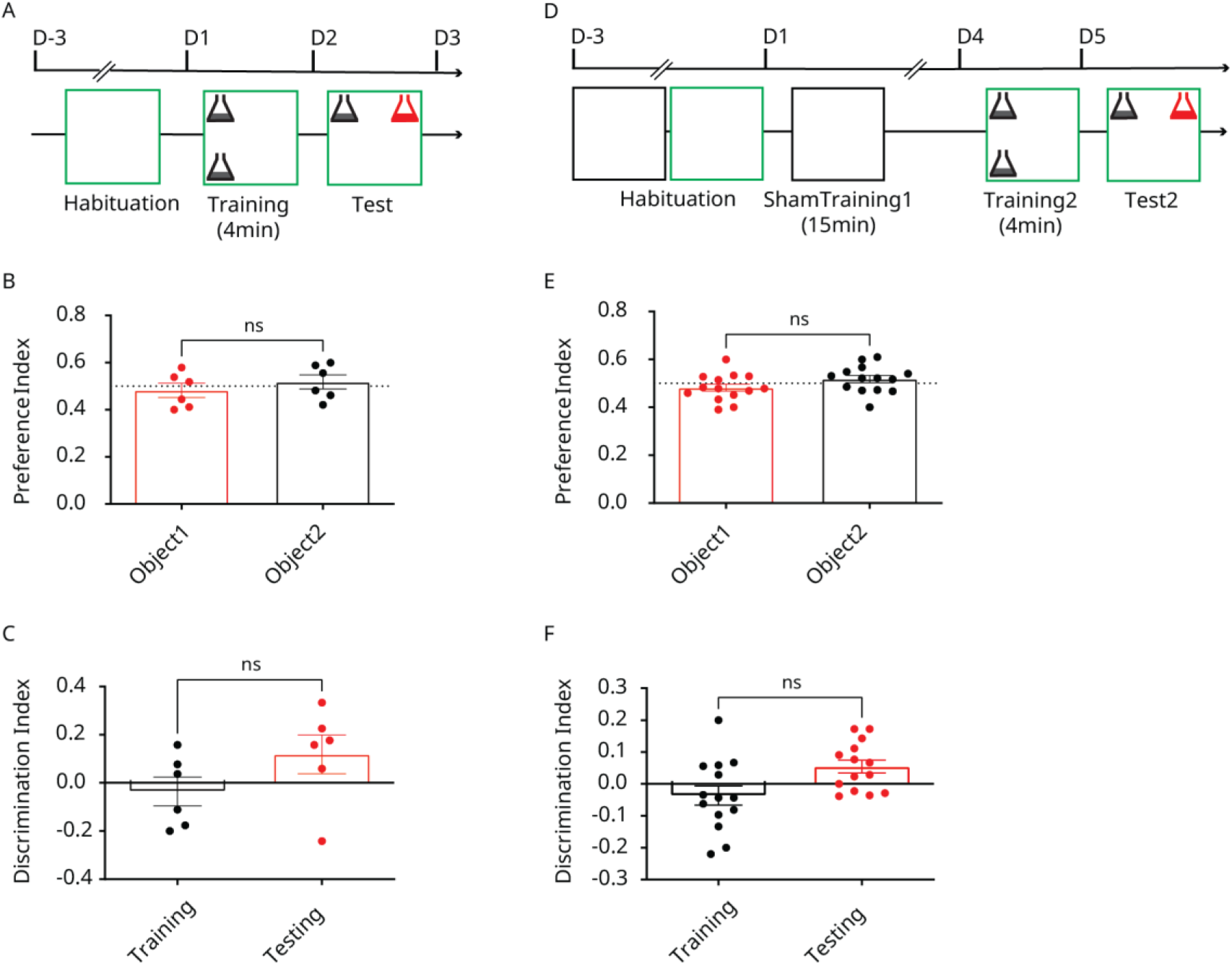
**(A)** Behavioral timeline for a single four-minute training episode in the modified context. **(B)** The preference index for both objects during training. **(C)** Discrimination of the novel location object during the test. *n* = 6 mice. **(D-F)** Same as **(A-C)** but for mice exposed first to one empty training episode in the original context, followed by a single four-minute training episode in the modified context. *n* = 14 mice. All statistical comparisons were performed using a paired Student’s t-test. ns, non-significant. Data are represented as mean ± standard error of the mean (SEM).

**Extended Data Figure3 (Related to Fig.2).**
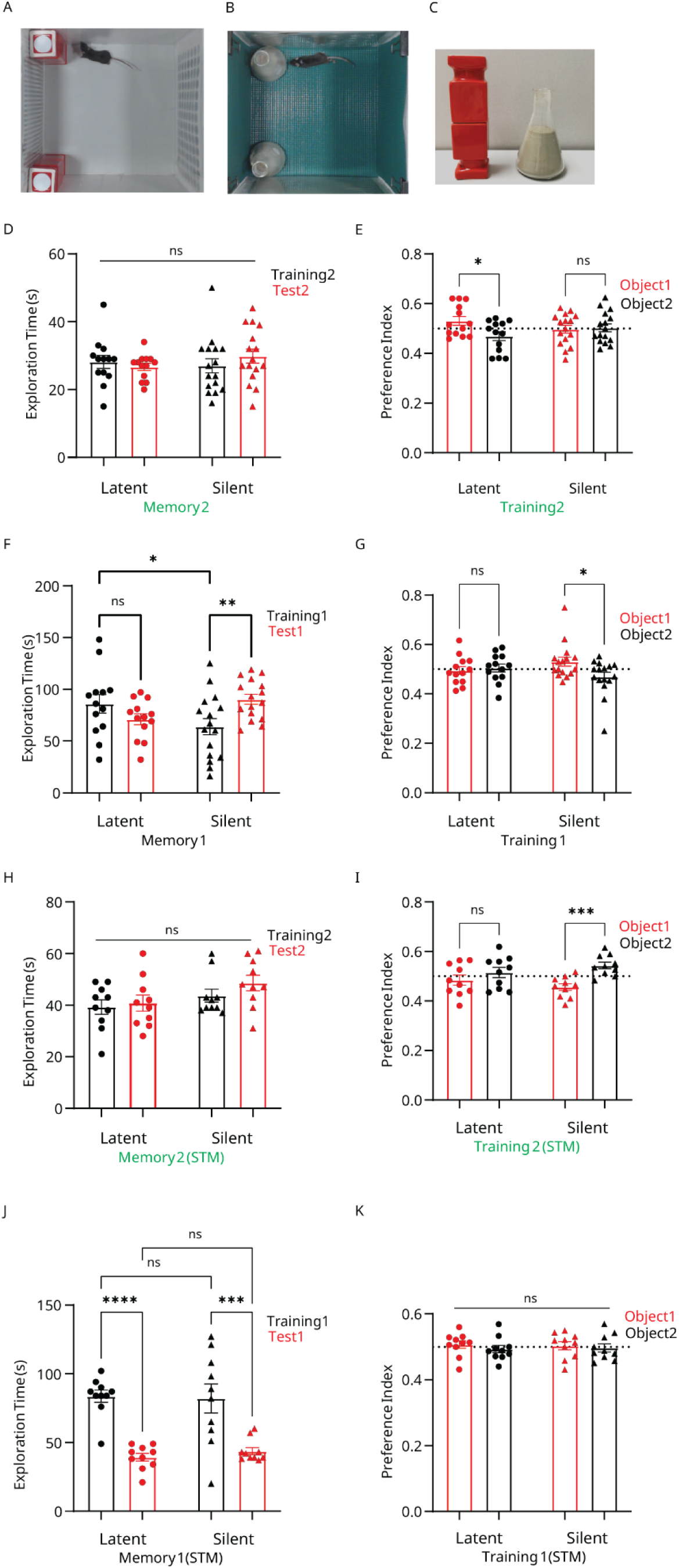
**(A)** Context and objects for the original memory. **(B)** Context and objects for the modified event. **(C)** and a side-by-side comparison for both objects. **(D)** Exploration times during training and testing of the modified memory for mice in Fig.2A. **(E)** The preference index for both objects during the modified memory training. **(F)** and **(G)** Similar to **(D)** and **(E),** but for the original memory. *n* = 13 and 16 mice for the latent and silent groups, respectively. **(H-K)** Similar to **(D-G)** but for the short-term memory task in Fig.2C. *n* = 10 mice/group. All statistical comparisons were performed using repeated measures two-way ANOVA. ns, non-significant; *, *P* < 0.05; ***, *P* < 0.001; ****, *P* < 0.0001. Data are represented as mean ± standard error of the mean (SEM).

**Extended Data Figure4 (Related to Fig.2).**
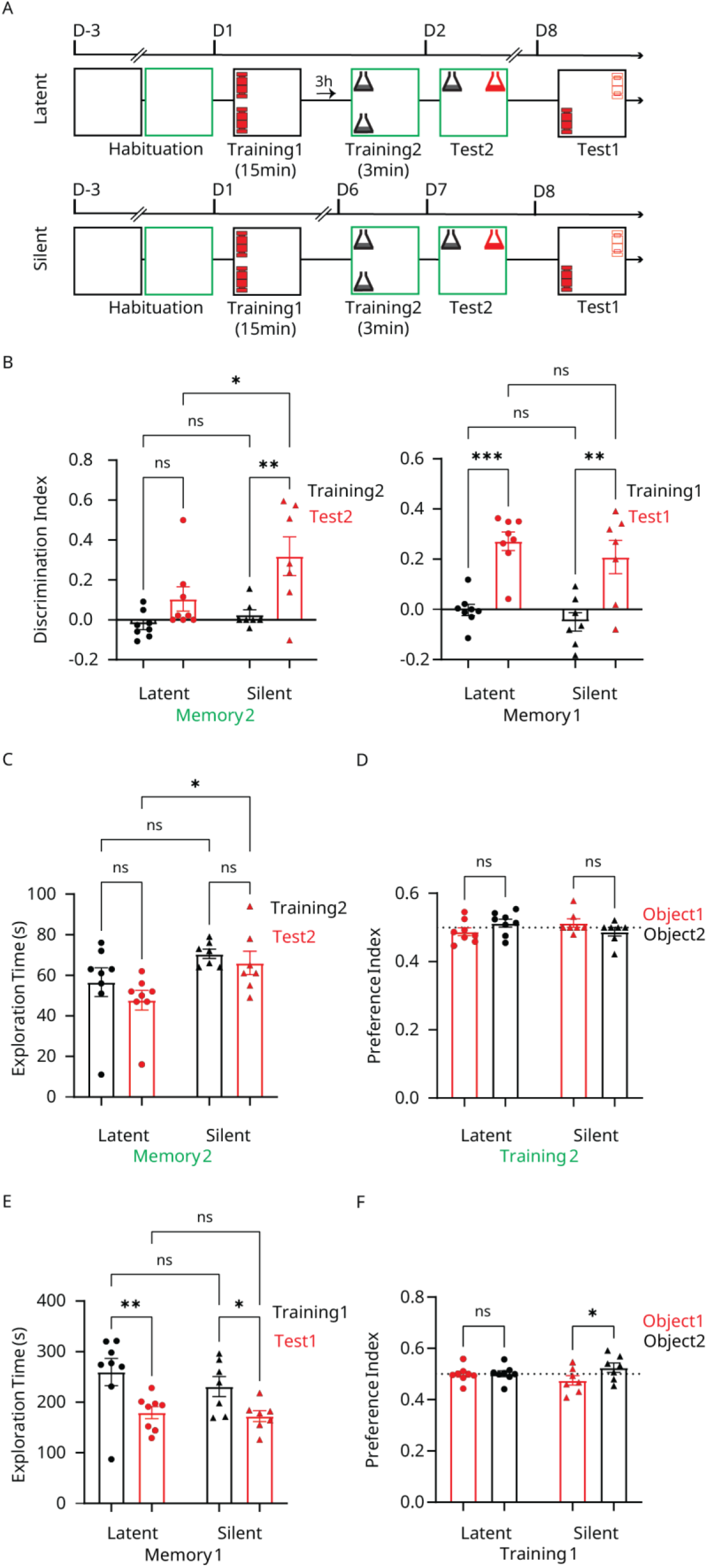
**(A)** Behavioral timeline for the latent (three hours after encoding, top) and silent (five days post encoding, bottom) NOPR engrams. **(B)** Discrimination of the novel object’s location at the modified (left) and original (right) memory tests for the latent and silent groups. **(C)** Exploration times during training and testing of the modified memory. **(D)** The preference index for both objects during the modified memory training. **(E) and (F)** Similar to **(C)** and **(D),** but for the original memory. *n* = 8 and 7 mice for the latent and silent groups, respectively. All statistical comparisons were performed using repeated measures two-way ANOVA. ns, non-significant; *, *P* < 0.05; **, *P* < 0.01; ***, *P* < 0.001. Data are represented as mean ± standard error of the mean (SEM).

**Extended Data Figure5 (Related to Fig.3).**
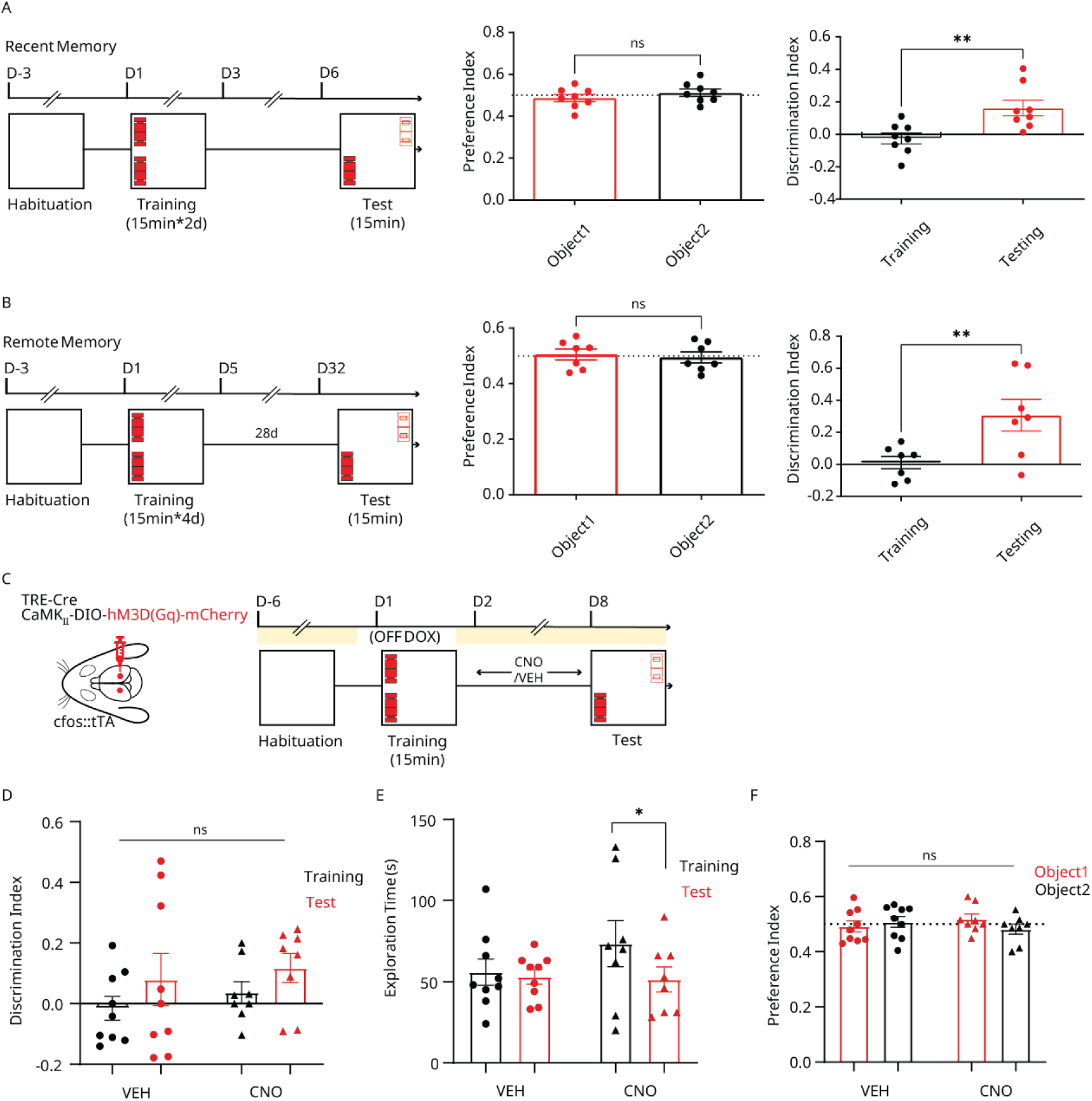
**(A)** Left, behavioral timeline to check for memory persistence at recent timepoints with a single retraining. Centre, preference index for both objects during training. Right, discrimination index for the novel location object during test. *n* = 8 mice. (**B**) Same as **A,** but for the remote memory test after four retraining epochs. *n* = 7 mice. **(C)** Design and behavioral timeline for the chemo-genetic maintenance of the NOPR engram activity. **(D)** Discrimination of the novel object’s location at the test for the VEH and CNO groups. **(E)** Exploration times during training and testing for the VEH and CNO groups. **(F)** The preference index for both objects during training. *n* = 9 and 8 mice for VEH and CNO, respectively. All statistical comparisons were performed using paired Student’s t-test (A, B) or repeated measures two-way ANOVA (C-F). CNO, clozapine-N-oxide; VEH, vehicle. ns: non-significant; **, *P* < 0.01. Data are represented as mean ± standard error of the mean (SEM).

**Extended Data Figure6 (Related to Fig.3).**
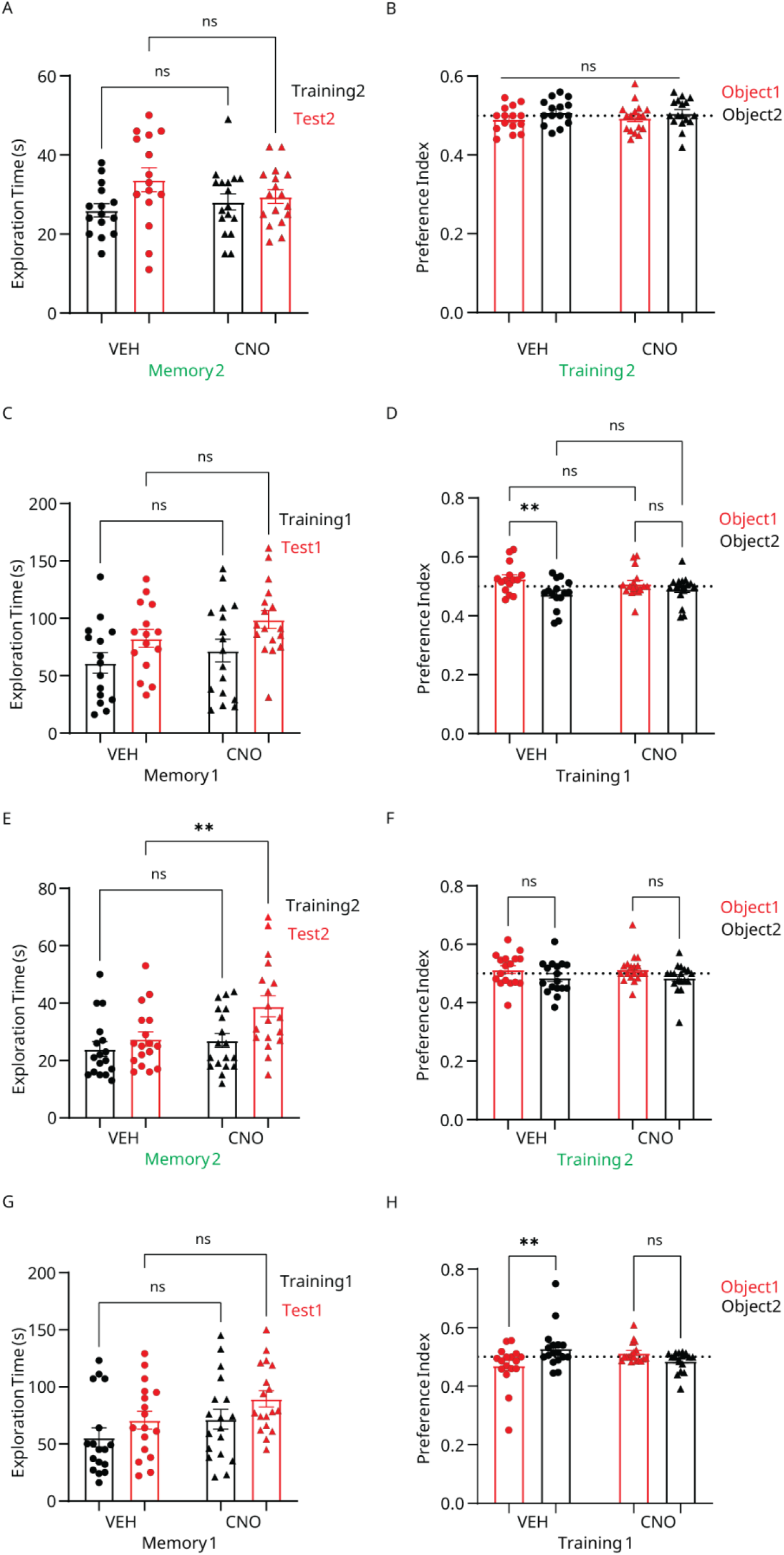
**(A)** Exploration times during training and testing of the modified memory for mice in Fig.3A. **(B)** The preference index for both objects during the modified memory training. **(C)** and **(D)** Similar to **(A)** and **(B),** but for the original memory. *n* = 15 and 17 mice for VEH and CNO, respectively. **(E-H)** Similar to **(A-D)** but for the engram inhibition experiment in Fig.3D. *n* = 17 and 18 mice for VEH and CNO, respectively. All statistical comparisons were performed using repeated measures two-way ANOVA. CNO, clozapine-N-oxide; VEH, vehicle. ns, non-significant; **, *P* < 0.01. Data are represented as mean ± standard error of the mean (SEM).

**Extended Data Figure7 (Related to Fig.4).**
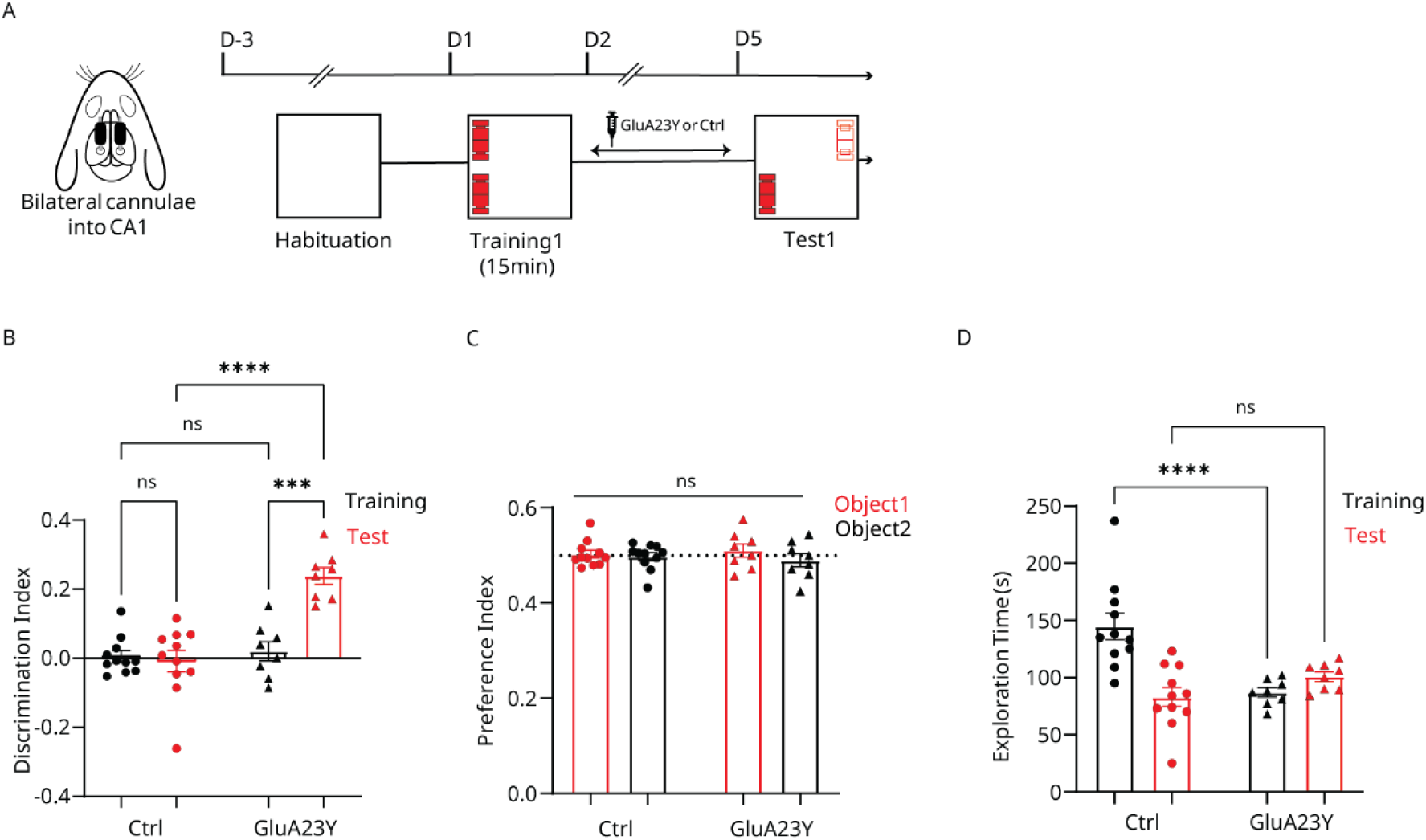
**(A)** Design and behavioral timeline for the maintenance of the NOPR engram activity through modulation of the synaptic localization of GluA_2_-containing AMPARs. **(B)** Discrimination of the novel object’s location at the test for the Ctrl and GluA23Y groups. **(C)** Exploration times during training and testing. **(D)** The preference index for both objects during training. *n* = 11 and 8 for the Ctrl and GluA23Y groups, respectively. All statistical comparisons were performed using repeated measures two-way ANOVA. Ctrl, control Tat-GluA2_3A_ peptide; GluA23Y, Tat-GluA2_3Y_ peptide. ns: non-significant; ***, *P* < 0.001; ****, *P* < 0.0001. Data are represented as mean ± standard error of the mean (SEM).

**Extended Data Figure8 (Related to Fig.4).**
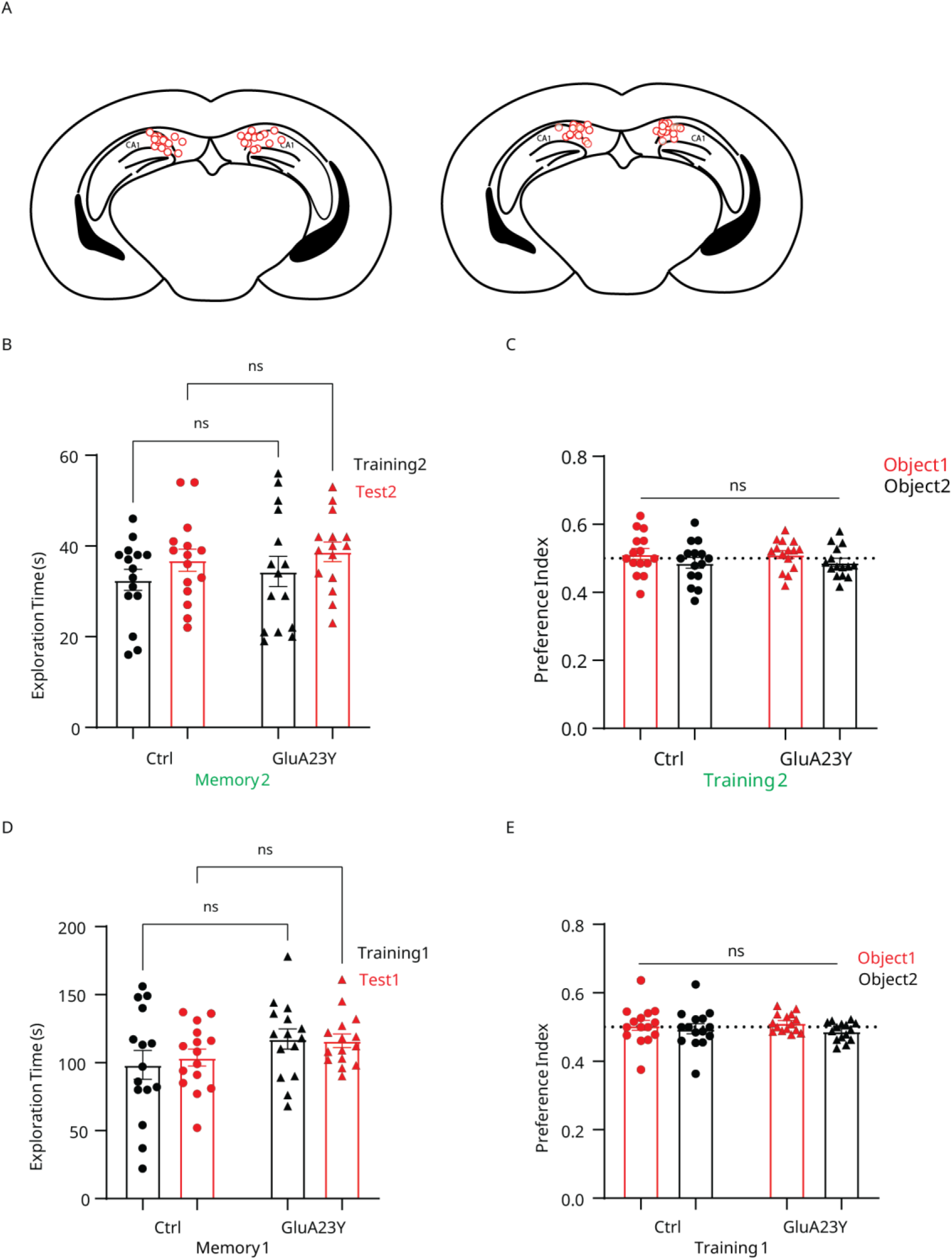
**(A)** Cannula placement for mice injected with Ctrl (left) or GluA23Y (right) peptides. **(B)** Exploration times during training and testing of the modified memory for mice in Fig.4A. **(C)** The preference index for both objects during the modified memory training. **(D)** and **(E)** Similar to **(B)** and **(C),** but for the original memory. *n* = 15 mice/group. All statistical comparisons were performed using repeated measures two-way ANOVA. Ctrl, control Tat-GluA2_3A_ peptide; GluA23Y, Tat-GluA2_3Y_ peptide. ns, non-significant. Data are represented as mean ± standard error of the mean (SEM).

**Extended Data Figure9 (Related to Fig.5).**
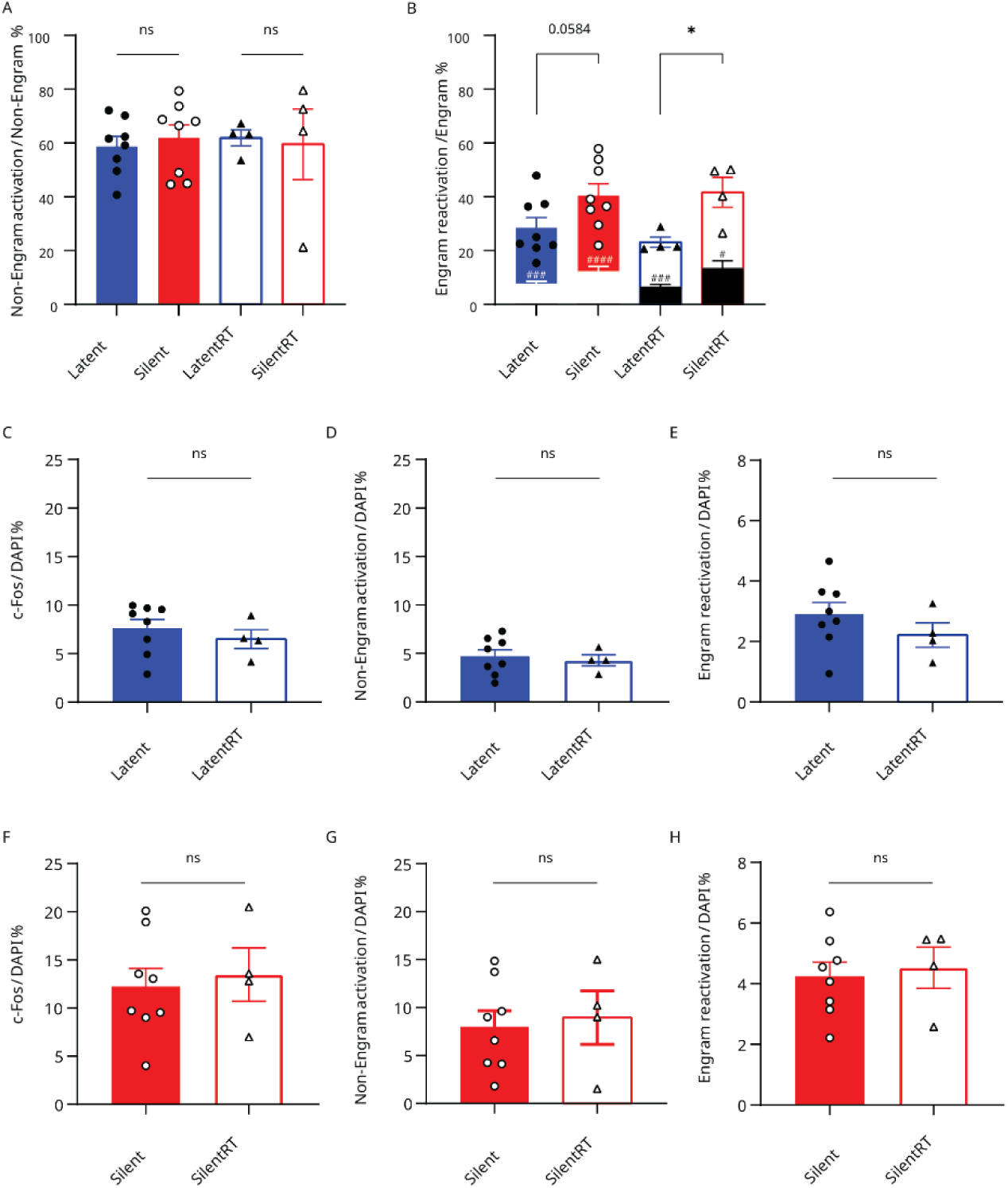
**(A)** Percentage of active non-engram cells (mCherry-negative, c-Fos-positive) within the total non-engram (mCherry-negative) population. **(B)** Percentage of reactivated engram (mCherry and c-Fos-positive) cells within the total engram (mCherry-positive) population. Inner columns represent the chance level for each group. **(C-E)** Within-state comparisons of the fraction of active neurons **(C)**, non-engram activation **(D)** and engram reactivation **(E)** for the Latent and Latent retraining groups. **(F-H)** Same as **(C-E)** but for the Silent and Silent retraining groups n = 8 and 4 mice per group for the modified and retraining events, respectively. All statistical comparisons across groups were performed using an unpaired Student’s t-test, while within-group comparisons against chance level were calculated with a paired Student’s t-test. ns, non-significant; *, P < 0.05 for group comparisons and #, P < 0.05; ###, P < 0.001; ####, P < 0.0001 for comparison to chance level. Data are represented as mean ± standard error of the mean (SEM).

**Extended Data Figure10 (Related to Fig.5).**
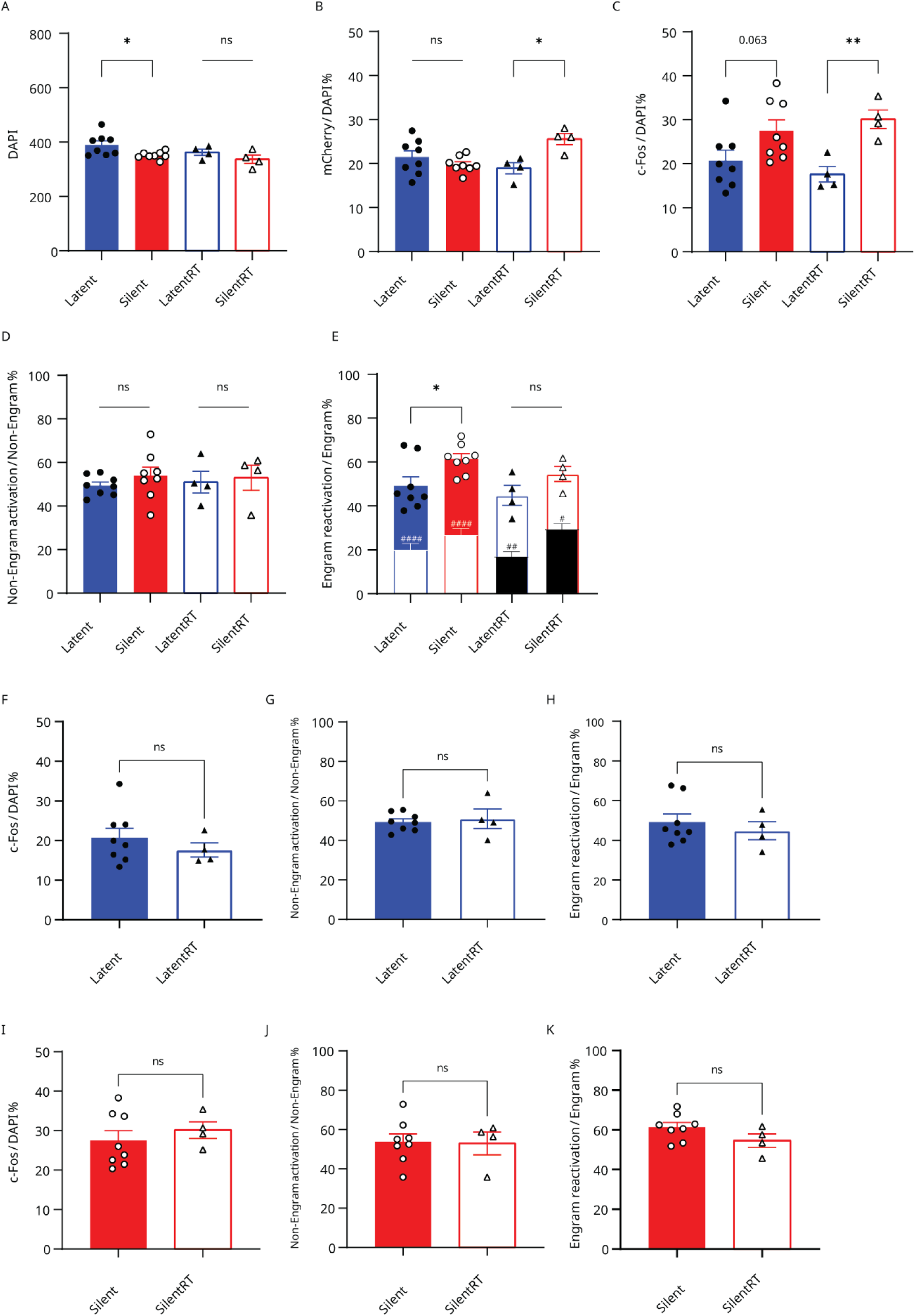
**(A)** DAPI-positive cells across groups. **(B)** Percentage of engram (mCherry-positive) cells from total counted cells across groups. **(C)** Percentage of active (c-Fos-positive) cells among the total counted cells across groups. **(D)** Percentage of active non-engram cells (mCherry-negative, c-Fos-positive) within the total non-engram (mCherry-negative) population. **(E)** Percentage of reactivated engram (mCherry and c-Fos-positive) cells within the total engram (mCherry-positive) population. Inner columns represent the chance level for each group. **(F-H)** Within-state comparisons of the fraction of active neurons **(F)**, non-engram activation **(G)**, and engram reactivation **(H)** for the Latent and Latent retraining groups. **(I-K)** Same as **(F-H)** but for the Silent and Silent retraining groups. *n* = 8 and 4 mice per group for the modified and retraining events respectively. All statistical comparisons across groups were performed using an unpaired Student’s t-test, while within-group comparisons against chance level were calculated with a paired Student’s t-test. ns, non-significant; *, P < 0.05 and **, P < 0.01 for group comparisons and #, P < 0.05; ##, P < 0.01; ####, P < 0.0001 for comparison to chance level. Data are represented as mean ± standard error of the mean (SEM).

## Notes

### Competing Interest Statement

The authors have declared no competing interest.

